# Thermodynamic Quality Control Reshapes Spatially Resolved Metabolic Flux Predictions in Prostate Cancer

**DOI:** 10.64898/2026.06.25.734531

**Authors:** Mohammad Reza Zargar, Sunayana Malla, Rajib Saha, Ratul Chowdhury

## Abstract

Spatial transcriptomics enables region-resolved investigation of tumor metabolism, yet translating gene expression into reliable metabolic flux predictions remains challenging because genome-scale models can admit thermodynamically infeasible cycles (TICs). Here, we integrated spatial transcriptomic data from twelve annotated regions of a single prostate tumor, spanning normal, inflamed, and Gleason 3+3 and 3+4 tissues, with Human-GEM using an E-Flux framework incorporating defined nutrient constraints, explicit cycle certification, and loopless flux variability analysis. An initial naïve implementation produced TICs in all twelve flux solutions, with total internal fluxes 190–890-fold higher than those obtained using the revised workflow. All flux solutions were certified free of TICs, and 94.7% of 17,592 loopless flux bounds were determined exactly. Within the analyzed tissue, predicted metabolic similarity was associated with spatial region (permutation p = 0.023) but not histological state (p = 0.52), revealing model-predicted heterogeneity even between regions sharing the same Gleason grade. Flux-based target prioritization identified fatty acid uptake-associated genes (CD36, FABPs, and SLC27A1/4), although the dominant uptake signal was confined to one malignant region, suggesting a localized metabolic hypothesis rather than an established therapeutic vulnerability. Analysis of the zinc–citrate–aconitase axis further demonstrated that zinc-dependent citrate redistribution was absent under baseline conditions and emerged only when oxidative demand, elimination of a cytosolic aconitase bypass, and strong mitochondrial aconitase inhibition coincided. These findings establish the importance of thermodynamic quality control and physiologically informed constraints in spatial metabolic modeling, while distinguishing computationally feasible flux predictions from experimentally validated metabolic mechanisms. The resulting framework provides a reproducible foundation for investigating spatial metabolic heterogeneity and prioritizing testable hypotheses for future experimental validation.

## 1. Introduction

Prostate cancer is the most frequently diagnosed solid malignancy among American men, with an estimated 299,010 new cases and 35,250 deaths in the United States in 2024 [1]. Its clinical course ranges from indolent disease suitable for surveillance to lethal, androgen independent metastatic disease [2]. Much of this variability is rooted in tumor heterogeneity. Primary prostate cancer is often multifocal, foci of different grade coexist within one gland, and genomic and transcriptional states vary across short distances, including in histologically benign tissue [3], [4]. Metabolism is one layer of this heterogeneity that is directly actionable, because metabolic enzymes and transporters can be targeted pharmacologically [5].

Prostate cancer metabolism differs from that of most solid tumors. Primary prostate tumors do not typically show the Warburg increase in glucose uptake and lactate secretion [6], [7]. Instead, they rely on lipid metabolism and on a citrate-centered metabolism inherited from normal prostatic epithelium. Normal glandular cells accumulate zinc, which inhibits the citrate → cis-aconitate reaction of mitochondrial aconitase (ACO2) and favors citrate secretion [8], [9]. Malignant epithelium loses ZIP1 (SLC39A1) zinc import and zinc, restores aconitase activity and oxidizes citrate [10], [11], [12], [13]. Prostate tumors also depend on lipids, through *de novo* synthesis, with FASN expression rising with Gleason grade [14], and through avid uptake of exogenous fatty acids, which CD36 inhibition suppresses in preclinical models [15]. Androgen-receptor (AR) signaling drives growth and lipid metabolism [16], [17]. Whether these programs vary across regions of the same tumor, and how such variation could inform treatment, is not known.

Spatial transcriptomics allows these questions to be asked *in situ* [18]. Berglund *et al*.[3] profiled prostate cancer tissue and annotated regions of normal, inflamed and malignant tissue, revealing transcriptional heterogeneity beyond histological boundaries. Genome-scale metabolic models (GEMs) provide a way to translate such expression data into metabolic phenotypes [19], [20], [21], [22]. They have been used to propose cancer-selective vulnerabilities [23], [24], [25], [26], [27], [28], [29], [30], typically with Human-GEM [31]. Wang *et al*.[32] built mCADRE-pruned, region-specific models from three of these sections and identified vulnerabilities *by in silico* gene knockout, including *de novo* lipogenesis genes such as FASN, ACLY and SLC25A1. Pruning represents each reaction present or absent, so flux magnitudes cannot be compared between regions, and that study did not report checks for thermodynamically infeasible cycles. Continuous bound scaling with E-Flux [33] instead keeps the full network and allows flux magnitudes to be compared between regions. Unlike Wang *et al*., our contribution is not simply another spatial metabolic reconstruction; it is a comparison of modeling assumption, flux-cycle certification, and how these choices alter downstream predictions.

Continuous flux predictions, however, are only as reliable as their implementation. FBA solutions can contain thermodynamically infeasible cycles (TICs), internal loops that carry flux without any net change of metabolites and violate the loop law [34], [35]. TICs inflate fluxes and distort flux variability analysis (FVA) ranges [36], and loopless FVA on a genome-scale human model is computationally demanding and rarely reported. Systematic comparisons have also shown that the method and its settings strongly affect predictions [37], [38]. Finally, regulation that is not reflected in transcript abundance, such as metal-dependent enzyme inhibition, cannot be captured. If such issues are not controlled, apparent spatial heterogeneity and candidate targets may be artifacts.

Despite growing interest in spatially resolved metabolic modeling, the reliability of flux predictions remains limited by several methodological challenges. Transcript abundance provides only an indirect estimate of enzymatic capacity, while assumptions concerning nutrient availability, reaction reversibility, biomass composition, and flux optimization can substantially influence predicted metabolic states. Moreover, conventional flux balance analysis may permit thermodynamically infeasible internal cycles, producing flux distributions that satisfy mass balance but cannot represent physically realizable steady-state metabolism. Although established methods exist to eliminate such cycles, their systematic application and verification in spatially resolved human genome-scale models remain important practical challenges. A further distinction is necessary: eliminating infeasible cycles improves mathematical consistency but does not independently establish physiological accuracy.

Here, we investigate these challenges using twelve spatially annotated regions from a single prostate tumor as a case study. We implement a quality-controlled Human-GEM workflow that combines expression-constrained reaction capacities, defined nutrient availability, explicit cycle certification, and loopless flux variability analysis with documented solution exactness. We first quantify how conventional modeling choices alter predicted flux distributions and candidate metabolic targets. Without these controls, every flux solution contained thermodynamically infeasible cycles, and open exchange reactions allowed the network to import lactate, citrate and pyruvate, which supported most of the predicted growth and led to *de novo* lipogenesis genes (FASN, HMGCR and SLC25A1) being prioritized as targets. With the revised workflow, every solution is certified free of internal cycles, growth is supported only by the defined medium, and lipid demand is met by uptake: FASN and ACLY carry no flux in any region, and cholesterol synthesis through HMGCR increases only when serum lipids are removed from the medium. These fluxes are defensible because the loop certificate is a linear program that is zero only when no internal cycle exists, and cycles are removed with all exchange and biomass fluxes held fixed, so no input or output of the model changes. We then examine whether models built with the revised workflow generate spatially structured metabolic differences and biologically testable hypotheses concerning lipid utilization and zinc-dependent citrate metabolism. Because the analysis is limited to one patient and lacks direct metabolic flux measurements, we interpret these biological results as exploratory predictions rather than validated disease mechanisms, and TCGA-PRAD [39] and DepMap [40] data serve as orthogonal checks rather than validation. The primary contribution is a reproducible framework for assessing the feasibility, sensitivity, and biological interpretability of spatially resolved metabolic models.

## 2. Materials and Methods

### 2.1 Spatial transcriptomics data

We used the spatial transcriptomics dataset of Berglund *et al*.[3] (CNGB STOmics STDS0000053), which comprises twelve annotated positions (P1.1–P4.3) in four regions (P1–P4) of tissue sections from a single prostate cancer patient (spot diameter 100 µm, center-to-center spacing 200 µm). Tissue states were taken from the original annotation: Normal (n = 6 positions: P1.1, P2.1, P3.2, P4.1, P4.2, P4.3), Inflammation (n = 2: P2.3, P3.1), Cancer Gleason 3+3 (n = 2: P1.2, P1.3) and Cancer Gleason 3+4 (n = 2: P2.4, P3.3). For each position, the mean expression across all spots in the annotated region was used; this averages over the cell types present (Section 4.4). Gene coverage in Human-GEM ranged from 28.7% (P4.3) to 67.0% (P2.1) **(Table 1).**

**Table 1.** The twelve spatial positions; tissue state, number of spots, genes detected, mean and maximum expression, gene coverage in Human-GEM (% of model genes detected above zero) and predicted biomass flux of the loop-free models (arbitrary units, a.u.)

| Position | Tissue state | Spots (n) | Genes detected | Mean expression | Max expression | Gene coverage (%) | Biomass (a.u.) |
| --- | --- | --- | --- | --- | --- | --- | --- |
| P1.1 | Normal | 432 | 8,696 | 0.283 | 70.0 | 51.4 | 0.0193 |
| P2.1 | Normal | 547 | 11,646 | 0.706 | 149.0 | 67.0 | 0.0326 |
| P3.2 | Normal | 523 | 6,338 | 0.101 | 7.5 | 39.9 | 0.0401 |
| P4.1 | Normal | 688 | 10,027 | 0.195 | 44.1 | 58.6 | 0.0069 |
| P4.2 | Normal | 324 | 9,513 | 0.420 | 58.0 | 57.0 | 0.0299 |
| P4.3 | Normal | 302 | 4,539 | 0.140 | 12.7 | 28.7 | 0.0381 |
| P2.3 | Inflammation | 483 | 8,282 | 0.164 | 16.5 | 49.9 | 0.0303 |
| P3.1 | Inflammation | 627 | 9,774 | 0.234 | 44.1 | 57.8 | 0.0152 |
| P1.2 | Cancer Gs3+3 | 406 | 11,285 | 0.732 | 393.9 | 65.3 | 0.0155 |
| P1.3 | Cancer Gs3+3 | 629 | 11,320 | 0.349 | 82.3 | 65.2 | 0.0276 |
| P2.4 | Cancer Gs3+4 | 448 | 10,064 | 0.293 | 54.4 | 58.6 | 0.0318 |
| P3.3 | Cancer Gs3+4 | 501 | 10,296 | 0.379 | 120.4 | 60.5 | 0.0144 |

### 2.2 Genome-scale metabolic model and E-Flux implementation

Human-GEM (12,931 reactions) was loaded with COBRApy [41]. A prostate cell biomass reaction (60 components; Table S2) and two transport reactions it requires were added (12,934 reactions), and the two generic biomass reactions were blocked. Uptake was restricted to a defined medium of RPMI-1640 plus the lipid, lipoprotein-cholesterol and trace-element components of 10% fetal bovine serum (60 uptake exchanges; Table S1); all other exchanges were secretion-only. For each position, GPR rules were evaluated with *AND* = *minimum* and *OR* = *sum* [33]; undetected genes took the position’s minimum positive expression; and capacities were set to *c_i_* = 1000 · *w_i_* / *max_k_ w_k_*, applied symmetrically to reversible reactions. Fluxes are therefore relative capacities in arbitrary units (a.u.), comparable across positions but not calibrated rates. Table 2 lists how these choices differ from the initial, conventional implementation.

**Table 2.** Implementation choices in the initial (conventional) E-Flux implementation and in the revised implementation. The right-hand column can serve as a checklist for spatial flux studies.

| Aspect | Initial implementation | Revised implementation |
| --- | --- | --- |
| Exchange reactions | All 1,660 open for uptake (lactate, citrate, pyruvate could be imported) | Defined RPMI-1640 + 10% FBS medium (60 uptake exchanges); all others secretion-only |
| Reversible reactions | Upper bound scaled; lower bound left at -1,000 | Symmetric bounds $[-c_i, c_i]$ |
| Isozymes (OR) / subunits (AND) | Maximum / minimum | Sum / minimum (arithmetic-mean AND as sensitivity) |
| Capacity normalization | $e_i/e_{min} \times 1,000$ (dataset-wide minimum) | $c_i = 1,000 \cdot w_i/\max_k w_k$ (per position) |
| Flux solution | One FBA vertex | pFBA, then CycleFreeFlux with exchanges and biomass fixed |
| Thermodynamic check | None | Loop certificate for every solution (all = 0) |
| TICs | Not assessed | 2,273 TIC-participating reactions, list certified complete |
| Flux variability | Standard FVA | Loopless FVA with exactness status for each bound |
| Flux units | Reported as mmol gDW<sup>-1</sup> h<sup>-1</sup> | Arbitrary units (relative capacities) |
| Reaction identifiers | Nine reported IDs pointed to other reactions | Verified against Human-GEM (Tables S3, S6) |
| TCGA comparison | GEPIA2 summary values; ZIP1 under a wrong Ensembl ID | Per-sample GDC TPM, Mann-Whitney with BH; ZIP1 not used |
| Spatial statistics | Mann-Whitney on pairwise distances | Position-level label permutation, $p = (b + 1)/(m + 1)$ |

### 2.3 Certified loop-free flux solutions and flux ranges

Linear programs were solved with HiGHS *[42]*. For each position, biomass was maximized, parsimonious FBA (pFBA) minimized total flux at that optimum *[43]*, and CycleFreeFlux removed any remaining cyclic flux with biomass and exchanges fixed *[44]*. Each solution was tested with a loop certificate, a linear program that is zero if and only if the flux vector contains no internal cycle *[35]*. Reactions that can participate in a TIC (TIC-participating reactions) of Human-GEM were enumerated with closed boundaries, and the list was certified complete by an additional linear program. Loopless FVA was computed at ≥ 95% of maximal biomass for 733 reactions. Each bound was labeled by how it was obtained (exact for acyclic directions, exact by certification or loop-breaking, exact by cycle-cut MILP, or conservative but backed by a loop-free flux vector). Formulations and algorithms are given in Supplementary Methods S1–S3. For comparison, one FBA solution per position from the initial implementation (Table 2) was audited with the same loop certificate (Tables S7-S9)

Throughout this study, thermodynamic consistency refers specifically to the absence of thermodynamically infeasible internal flux cycles. The loop certificate establishes that each reported flux solution contains no such cycle under the specified network constraints. This does not constitute a concentration-dependent thermodynamic flux analysis, nor does it establish reaction feasibility from experimentally measured metabolite chemical potentials or Gibbs free energies. Accordingly, loop-free solutions are interpreted as mathematically admissible steady-state flux distributions rather than uniquely determined physiological flux states.

### 2.4 Spatial heterogeneity

Pathway activity was computed as the sum of the absolute fluxes (Z|*v*|) over the reactions of each Human-GEM subsystem. Metabolic distance between positions was 1 − *Pearson r* over all 12,934 fluxes. Because each position contributes to several pairs, region and state effects were tested by permuting labels at the position level (10,000 permutations; Mann–Whitney U of same versus different label distances as statistic; *p* = (*b* + 1)/(*m* + 1) [45]). Heterogeneity within a state was summarized by the distance between positions sharing a diagnosis. Biomass was compared across states by the Kruskal–Wallis test [46]. Further details, including principal component analysis [47], are in Supplementary Methods S5.

### 2.5 Androgen-receptor activity

AR activity was scored per position as the mean expression of AR, KLK3, KLK2, TMPRSS2, FKBP5, NKX3-1 and STEAP2 [17], [48]. Spearman correlations with the 60 active pathway activities were adjusted by Benjamini–Hochberg [49] and are exploratory (n = 12).

### 2.6 Target scoring and lipid dependence

Gene-linked reactions were scored as *Cancer*_*flux* × *max*(log_2_ *FC*, 0) × (1 + *ΔFVA_norm_*), where *Cancer*_*flux* is the mean |*v*| of the four cancer positions, log_2_ *FC* compares cancer with the six normal positions, and *ΔFVA_norm_* is the normalized gain of loopless FVA span in cancer. Gene scores were the maximum over each gene’s reactions; genes with a score ≥ the 85th percentile, Cancer_flux ≥ the median and log_2_ *FC*≥ 0.5 were prioritized. The ranking was recomputed with standard FVA, without the ΔFVA term and with an additive rank-sum score (Supplementary Methods S4). To test lipid dependence, all models were re-solved without uptake of palmitate, oleate, stearate, arachidonate and cholesterol (Supplementary Methods S4).

### 2.7 Energy metabolism and the zinc-citrate axis

LDH (MAR04388) flux and its loopless range at 95% and 100% of maximal biomass were computed, together with lactate mass balance and ATP production by source. A loopless LDH minimum ≤ 0 shows only that growth does not require LDH in the model. For the zinc–citrate axis we examined (a) spatial expression of axis genes and ACO2 in TCGA-PRAD per-sample data, and (b) a test whose design, outcomes and decision rule were fixed before any simulation (Supplementary Note S1). In the test, the citrate → cis-aconitate capacity of ACO2 [8] was scaled by 1/(1 + *θz*), with relative zinc z = 1 (normal) or 0.31 (cancer; 62–75% less zinc in malignant tissue [9]) and θ ∈ {0, 1, 3, 10, 30}. Every position was simulated under both zinc levels, with two AND rules. After this test, a post hoc analysis examined when zinc could affect flux at all (Supplementary Note S2). Uptake of the medium carbon sources was capped in proportion to their RPMI-1640 concentrations, at three levels relative to the unconstrained glucose uptake (×1, ×0.5, ×0.25); the cytosolic aconitase route (ACO1, with citrate export and isocitrate return) was left open or blocked; θ ∈ {0, 10, 30, 100, 300}; and m-aconitase was also blocked completely to test whether it is required.

### 2.8 External datasets

TCGA-PRAD expression (GDC STAR-TPM; 497 primary tumors, 51 solid-tissue normals) was compared by two-sided Mann–Whitney tests with Benjamini–Hochberg adjustment. DepMap 24Q4 Chronos gene effects were taken for five prostate adenocarcinoma lines (PC-3, VCaP, 22Rv1, LNCaP, DU145) and, to assess selectivity, five non-cancerous prostate lines (WPE1-NA22, BPH-1, P4E6, Shmac 4, Shmac 5); an effect < −0.5 was considered essential. These data test bulk transcript abundance and in vitro essentiality, which differ from predicted fluxes.

## 3. Results

### 3.1 Modeling assumptions and infeasible cycles substantially alter predicted metabolic fluxes

We first evaluated the extent to which commonly used implementation choices influence spatially resolved genome-scale flux predictions. The initial, conventional E-Flux implementation (Table 2) generated solutions containing thermodynamically infeasible internal cycles in all twelve spatial positions (loop certificate 74–125; Figure 1a). Cycles carried 1.2–6.8% of internal flux (Figure S1b), and internal flux totals (48,722–66,322 a.u.) were 190- to 890-fold higher than those obtained using the revised workflow (57–310 a.u.). Sequential modification of reaction bounds, nutrient uptake constraints, and flux optimization procedures revealed that these differences were not attributable to cycle elimination alone (Figure 1b; Table S7). In particular, open uptake of every exchanged metabolite allowed the network to import lactate, citrate and pyruvate, which supported most of the predicted growth: mean biomass fell from 2.92 to 0.025 a.u. once uptake was restricted to the defined medium, demonstrating the importance of extracellular nutrient assumptions. Unscaled reverse bounds and a single FBA vertex left internal fluxes poorly determined: re-solving the same problem with HiGHS gave the same biomass but internal totals of 436–511,678 a.u. between positions and a cycle in only one position (Table S7), so whether a single FBA solution contains cycles depends on the solver and must be certified rather than assumed. In addition, nine reaction identifiers used to report key fluxes in the initial implementation pointed to unrelated reactions (Table S3).

**Figure 1.**
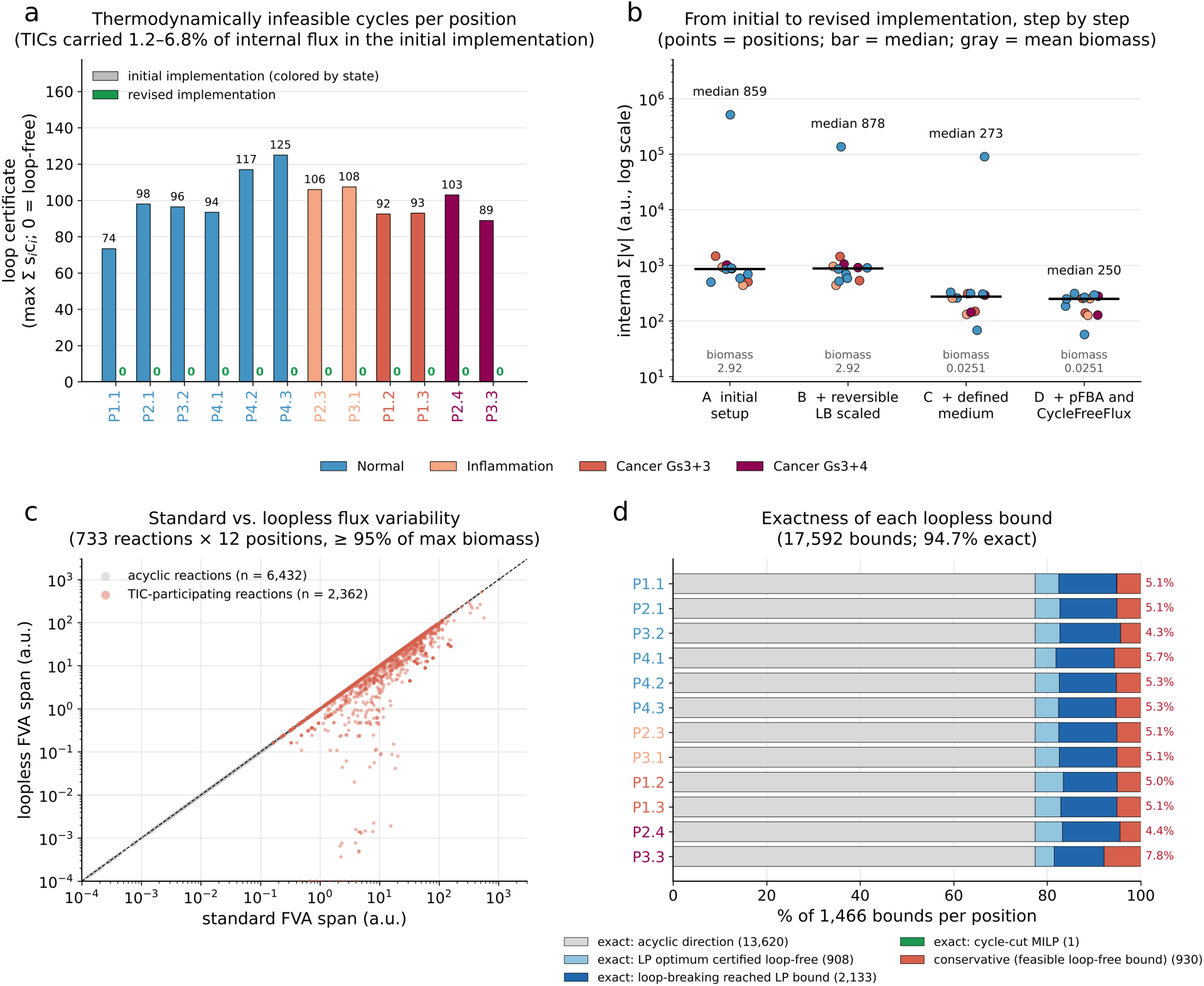
Certified loop-free flux predictions. **(a)** Loop certificate (maximal sign-conformal internal cycle flux; 0 = loop-free) of each position’s flux solution in the initial implementation (bars colored by tissue state) and in the revised implementation (green; 0 in all positions). **(b)** Internal Σ|v| as the initial setup is corrected one step at a time, all solved with HiGHS: A, open exchanges, upper-bound-only scaling and FBA; B, reverse bounds of reversible reactions scaled; C, defined RPMI-1640 + serum medium; D, pFBA followed by CycleFreeFlux. Points are positions, black bars are medians and gray numbers give the mean biomass (a.u.). **(c)** Standard versus loopless FVA span for 733 reactions in 12 positions; TIC-participating reactions in red, acyclic reactions (on the diagonal) in gray. Where a loopless bound is conservative, the true loopless span lies between the two values. **(d)** How each loopless bound was obtained; percentages at right are the conservative bounds per position.

The revised workflow generated flux solutions certified free of internal cycles across all twelve positions (Figure 1a). Of the 11,274 internal reactions of Human-GEM, 2,273 (20.2%) can participate in a TIC, mostly transport reactions (Figure S1a; Table S4), and a completeness test confirmed that none was missed. Of 17,592 loopless flux variability bounds (733 reactions in 12 positions), 94.7% were exact, while the remaining 5.3% (4.3–7.8% per position) represented conservative bounds supported by feasible loop-free solutions (Figure 1d). For TIC-participating reactions, loopless ranges were often more than an order of magnitude narrower than standard FVA ranges (Figure 1c), and across all reactions the loop law removed 4.0–20.1% of the total standard FVA span per position (Figure S1c); where bounds are conservative, these reductions are upper estimates. These results establish the computational feasibility of systematic cycle certification and uncertainty-aware flux range estimation in spatially resolved Human-GEM models. Importantly, they demonstrate that apparently large metabolic fluxes can arise from model implementation choices rather than differences in the underlying transcriptomic data. The resulting workflow provides a quality-control foundation for subsequent biological hypothesis generation, although independent measurements remain necessary to determine physiological accuracy. All results below are based on these loop-free solutions.

### 3.2 Predicted metabolic similarity is associated with spatial region in the analyzed tissue

Within the analyzed tissue, predicted flux profiles showed greater similarity among positions sharing a spatial region than among positions from different regions (mean correlation distance 0.029 versus 0.037; position-level permutation p = 0.023). In contrast, positions sharing a histological state were not significantly more similar than those with different annotations (0.038 versus 0.034; p = 0.52; Figure 2a, b). In the clustering, the two inflamed positions from different regions (P2.3, P3.1) had nearly identical predicted profiles, whereas cancer positions did not form a cluster, and the Gleason 3+3 position P1.2 and the normal position P1.1 were the two most distinct positions overall. Predicted biomass did not differ between states (Kruskal–Wallis p = 0.73; Figure S2b), and the first principal component of pathway activity mainly reflected overall flux scale (Figure S2a). These findings indicate spatial structure in the model-predicted flux distributions within this specimen. However, because the twelve positions originate from one patient and spatially adjacent positions may share both biological and technical characteristics, these results should not be interpreted as evidence that tissue location generally explains prostate cancer metabolism better than histological grade.

**Figure 2.**
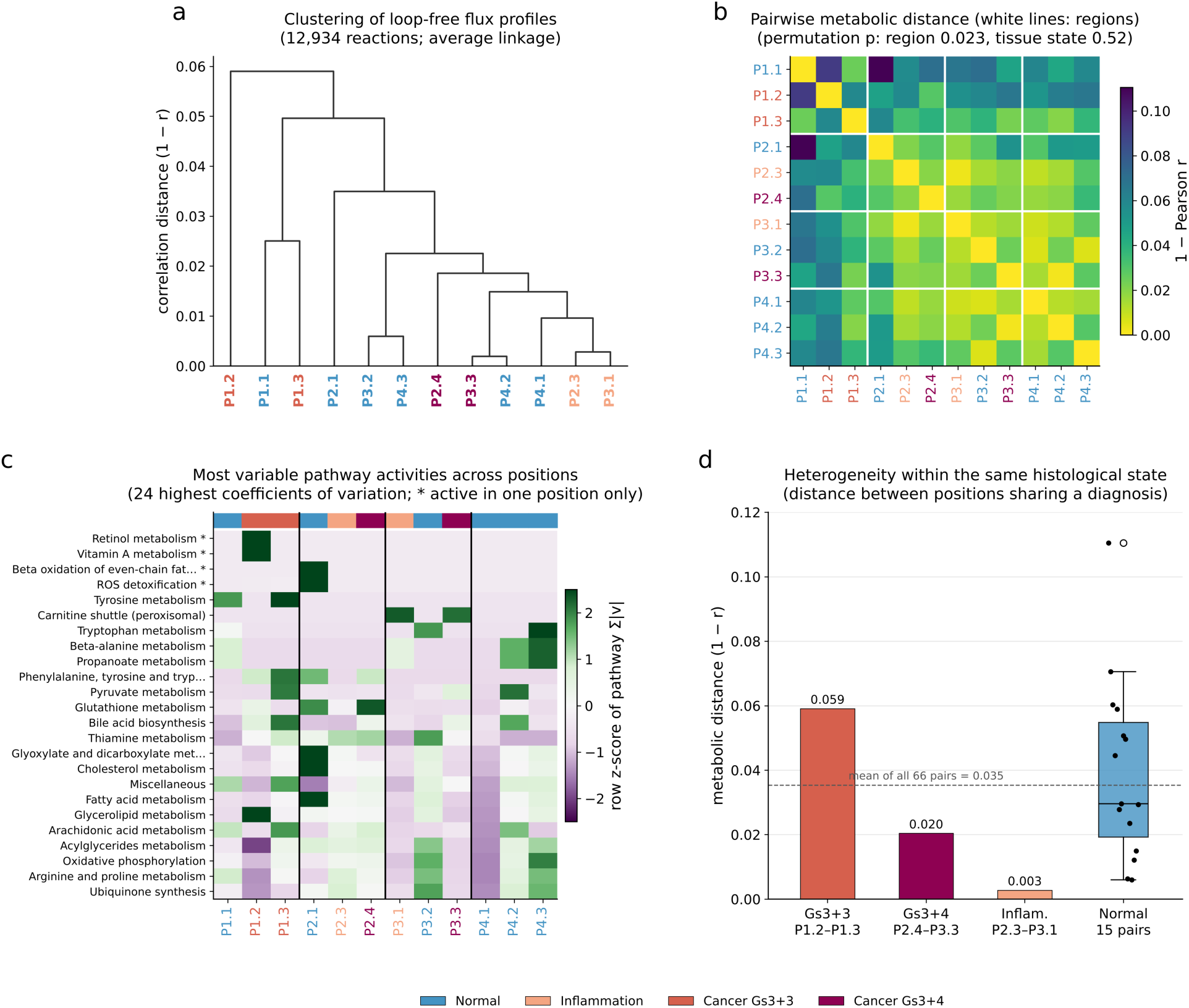
Spatial metabolic heterogeneity. **(a)** Average-linkage clustering of the twelve positions by correlation distance over 12,934 loop-free fluxes; labels are colored by tissue state. **(b)** Pairwise metabolic distance (1 − Pearson r), positions ordered by region; white lines separate regions P1–P4. Permutation p-values shuffle region or state labels at the position level (10,000 permutations). **(c)** The 24 pathway activities with the highest coefficient of variation across positions (row z-score), with positions ordered by region and the color bar showing tissue state; * marks pathways active in only one position. **(d)** Metabolic distance between positions sharing a diagnosis; the box shows the 15 pairs of normal positions and the dashed line the mean of all 66 pairs.

### 3.3 Exploratory within-grade comparisons reveal differences between annotated positions

Exploratory comparisons also revealed differences between positions assigned the same diagnosis. The two Gleason 3+3 positions (P1.2 and P1.3), which come from the same region, had a correlation distance of 0.059, exceeding the mean distance across all position pairs (0.035), whereas the two Gleason 3+4 positions (0.020) and the two inflamed positions (0.003) were similar (Figure 2d). Normal positions varied most (0.006–0.110 across 15 pairs), and the most distant pair in the tissue was two normal positions from different regions (P1.1 and P2.1). Pathway-level differences were often confined to single positions (Figure 2c): retinol and vitamin A metabolism were active only in P1.2, ROS detoxification and peroxisomal β-oxidation only in P2.1, and tryptophan metabolism in three normal positions. These observations illustrate that expression-constrained metabolic models can generate distinct predicted states within a shared histological annotation. Whether these differences reflect cell-intrinsic metabolic reprogramming, variation in tissue composition, or technical effects remains unresolved and requires evaluation in independent, multi-patient spatial datasets.

AR activity showed the same pattern (Figure 3a). It ranged from 1.1 (P3.1, Inflammation) to 79.5 (P1.2, Gleason 3+3). The second Gleason 3+3 position scored 16.1, the Gleason 3+4 positions 11.2 and 22.4, and one normal position (P2.1) 26.2. No pathway activity correlated with AR score after multiple-testing correction (strongest: ROS detoxification, ρ = 0.68, p = 0.015, q = 0.49; Figure 3b–e), and these pathways carried very small fluxes. At this resolution, AR activity therefore marks heterogeneity between regions but does not explain their flux differences. P1.2 combined the highest AR activity, the highest mean expression and the only fatty-acid uptake flux (Section 3.4), making it a candidate distinct metabolic niche within this tumor that would need confirmation in additional samples.

**Figure 3.**
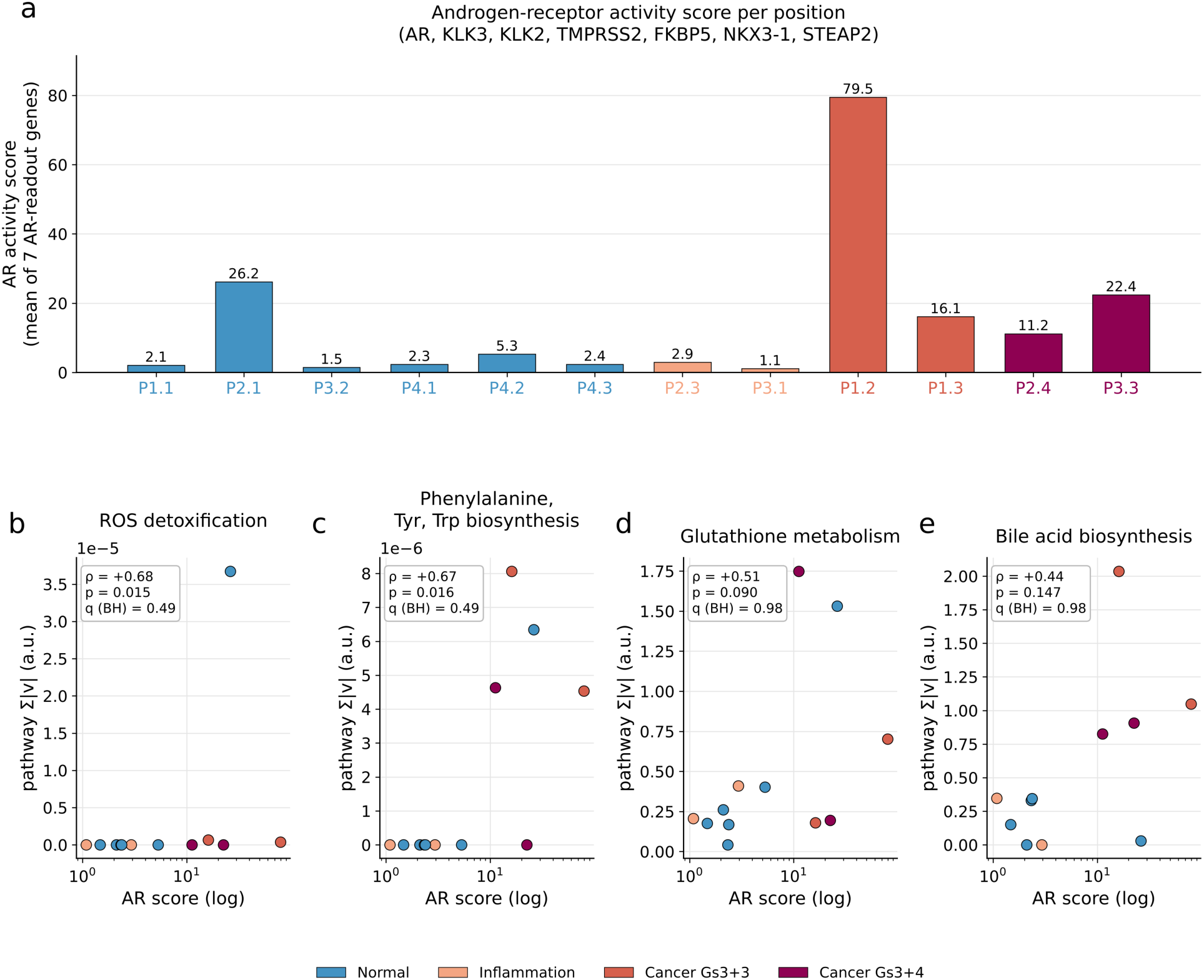
Androgen-receptor activity across positions. **(a)** AR activity score (mean expression of AR, KLK3, KLK2, TMPRSS2, FKBP5, NKX3-1 and STEAP2) per position, ordered by tissue state. **(b–e)** The four pathway activities with the smallest p-values against AR score. ρ, Spearman correlation; q, Benjamini–Hochberg-adjusted p over 60 pathways.

### 3.4 Flux-based prioritization identifies a region-specific fatty acid uptake hypothesis

Ninety-four gene-linked reactions carried higher mean flux in cancer than in normal positions. Thirteen genes, representing four reactions, met the prioritization criteria (Figure 4; Table 3). Seven of them share the top-ranked reaction, long-chain fatty-acid uptake (MAR00155: CD36, FABP2, FABP3, FABP4, FABP5, SLC27A1, SLC27A4; score 14.96), and therefore constitute one reaction-level signal rather than seven independently supported therapeutic targets. Four are monocarboxylate transporters (MAR06048: SLC16A3, SLC16A5, SLC16A7, SLC16A8; 2.83), and the others are CYP27A1 (1.58) and the mitochondrial dicarboxylate carrier SLC25A10 (1.25). SLC7A5 ranked 12th, below the prioritization threshold, and SLC25A1 ranked 52nd. FASN, ACLY, ACACA, HMGCR, PKM, HK2, ACO2, ACSL3 and SCD carried no higher flux in cancer than in normal positions and were not candidates.

**Figure 4.**
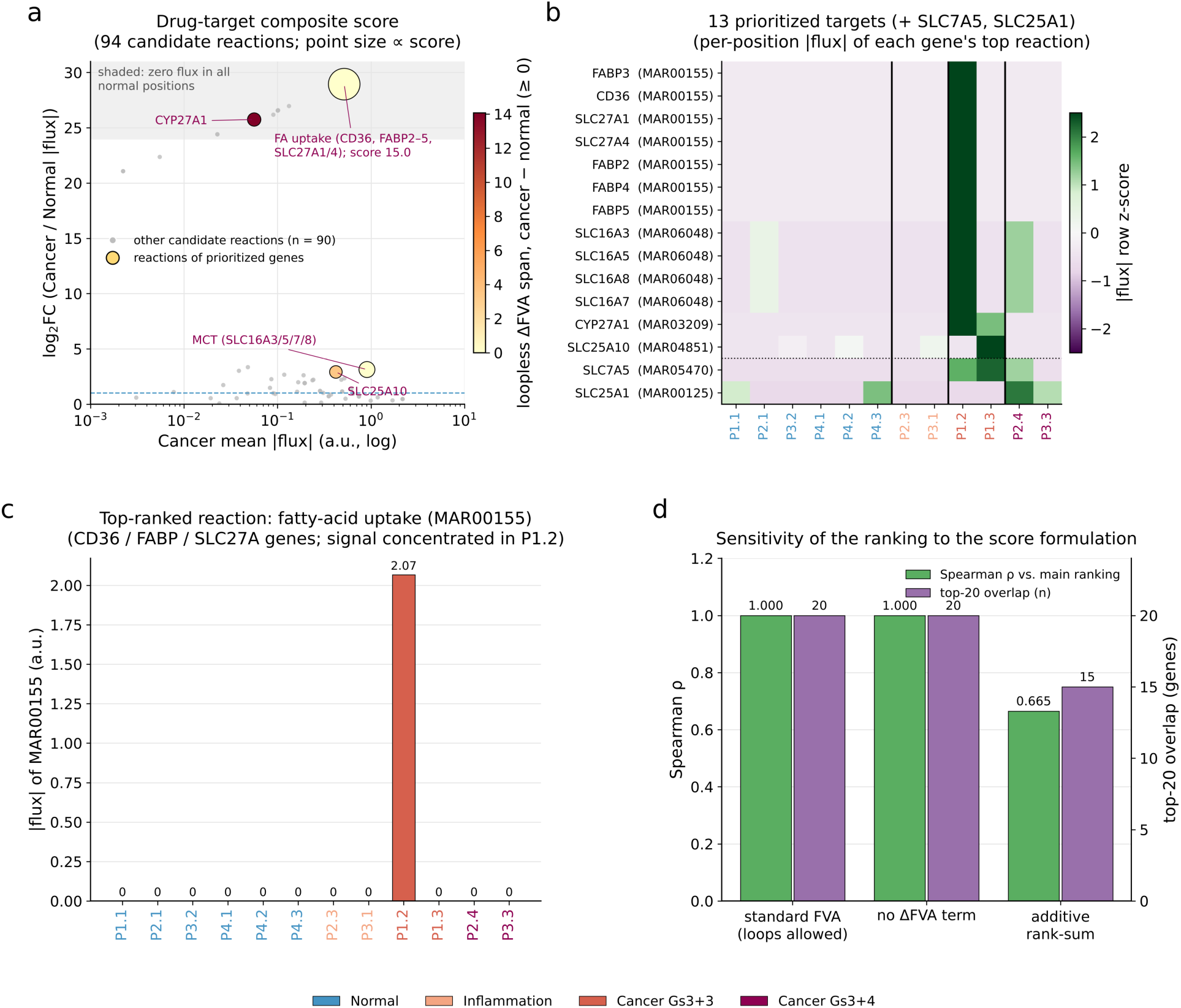
Flux-based target scoring. **(a)** Cancer mean |flux| versus log_2_FC (cancer/normal) for the 94 candidate reactions; point size is proportional to the score and color shows the loopless ΔFVA span (cancer − normal, clipped at 0). The shaded band contains reactions with zero flux in every normal position. **(b)** Per-position |flux| (row z-score) of the top reaction of each prioritized gene, plus SLC7A5 and SLC25A1 (below the dotted line); vertical lines separate tissue states. **(c)** Flux of the top-ranked reaction, fatty-acid uptake (MAR00155), in each position. **(d)** Sensitivity of the gene ranking to the score formulation: Spearman ρ with the main ranking and overlap of the top 20 genes.

**Table 3.** Prioritized targets and refringence genes: model rank and score, TCGA-PRAD differential expression and DepMap 24Q4 essentiality (mean Chronos effect in five prostate cancer and five non-cancerous prostate lines; percentage of cancer lines with effect < −0.5). “–”, not a model candidate or not available.

| Gene | Model rank | Score | TCGA $\log_2 \text{FC}$ | TCGA adj. p | DepMap, cancer | DepMap, non-cancer | % cancer lines essential | Prioritised |
| --- | --- | --- | --- | --- | --- | --- | --- | --- |
| FABP3 | 1 | 14.96 | - | - | 0.10 | -0.01 | 0 | yes |
| CD36 | 2 | 14.96 | -1.40 | $1.9 \times 10^{-6}$ | -0.13 | -0.11 | 20 | yes |
| SLC27A1 | 3 | 14.96 | - | - | -0.11 | -0.11 | 0 | yes |
| SLC27A4 | 4 | 14.96 | - | - | 0.00 | -0.04 | 0 | yes |
| FABP2 | 5 | 14.96 | - | - | 0.11 | 0.04 | 0 | yes |
| FABP4 | 6 | 14.96 | - | - | 0.08 | 0.05 | 0 | yes |
| FABP5 | 7 | 14.96 | -0.10 | 0.932 | - | - | - | yes |
| SLC16A3 | 8 | 2.83 | - | - | -0.08 | 0.03 | 0 | yes |
| SLC16A5 | 9 | 2.83 | -0.50 | 0.002 | -0.09 | -0.11 | 0 | yes |
| SLC16A8 | 10 | 2.83 | - | - | 0.01 | 0.01 | 0 | yes |
| SLC16A7 | 11 | 2.83 | - | - | -0.16 | -0.14 | 0 | yes |
| CYP27A1 | 14 | 1.58 | - | - | 0.07 | -0.01 | 0 | yes |
| SLC25A10 | 18 | 1.25 | - | - | -0.24 | -0.21 | 0 | yes |
| SLC7A5 | 12 | 2.35 | -0.99 | $9.3 \times 10^{-5}$ | -0.15 | -0.28 | 0 | no |
| SLC25A1 | 52 | 0.33 | +0.21 | $3.4 \times 10^{-4}$ | -0.25 | -0.17 | 20 | no |
| FASN | - | - | +1.65 | $3.3 \times 10^{-11}$ | -0.11 | -0.52 | 0 | no |
| ACLY | - | - | +0.38 | $5.1 \times 10^{-4}$ | -0.34 | 0.03 | 20 | no |
| ACSL3 | - | - | +0.26 | 0.049 | -0.27 | -0.21 | 20 | no |
| HMGCR | - | - | -0.41 | 0.075 | -1.25 | -1.24 | 100 | no |
| PKM | - | - | -0.36 | $1.6 \times 10^{-5}$ | -0.58 | -0.72 | 80 | no |
| HK2 | - | - | +0.57 | $1.9 \times 10^{-4}$ | -0.23 | -0.33 | 0 | no |
| ACO2 | - | - | -0.18 | 0.092 | -0.49 | -1.00 | 60 | no |
| ACACA | - | - | - | - | -0.24 | -0.71 | 0 | no |
| SCD | - | - | - | - | -0.62 | -1.01 | 60 | no |

The uptake signal was spatially restricted. Fatty-acid uptake flux was non-zero only in P1.2 (2.07 a.u.; Figure 4c), and monocarboxylate-transporter flux only in P1.2, P2.4 and P2.1 (Figure 4b). The ranking was insensitive to the FVA term: with standard instead of loopless FVA, or without the ΔFVA term, Spearman ρ was 1.00 and 0.9998, with 20 of the top 20 genes shared. It was moderately sensitive to the functional form of the score (additive rank-sum: ρ = 0.67, 15/20 shared; Figure 4d). When serum lipids were removed from the medium, biomass fell by 8–17% across states and HMGCR and ACACA flux rose (about 290-fold and 1.8-fold), but citrate synthase, ACLY and the FASN acetyl-transacylase step remained at zero (Figure 5c). Lipid supply in these models is therefore met by uptake whenever exogenous lipids are available.

**Figure 5.**
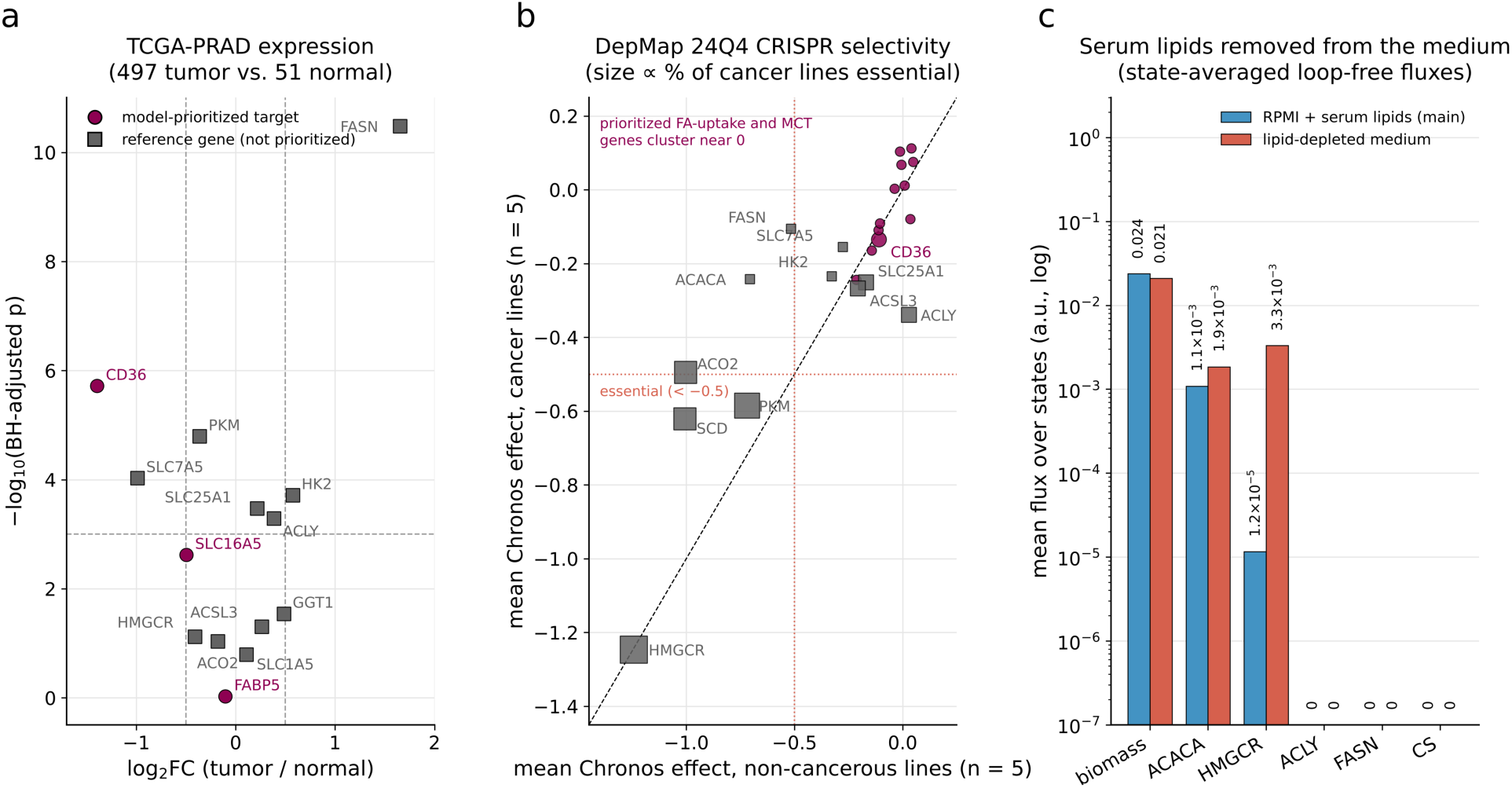
External datasets and lipid dependence. **(a)** TCGA-PRAD differential expression (497 tumors vs 51 normals; Mann–Whitney with Benjamini–Hochberg adjustment) of model-prioritized genes (circles) and reference genes that were not prioritized (squares; de novo lipogenesis and the citrate axis, glycolysis, and amino-acid transport and glutathione metabolism, several of them proposed as targets in prostate cancer [7]; dashed lines mark |log_2_FC| = 0.5 and adjusted p = 0.001. **(b)** DepMap 24Q4 Chronos effect in prostate cancer lines (n = 5) versus non-cancerous prostate lines (n = 5); points on the diagonal are not cancer-selective; size is proportional to the percentage of cancer lines in which the gene is essential (< −0.5). **(c)** State-averaged fluxes in the main medium and in a lipid-depleted medium.

The flux-based ranking prioritized genes associated with long-chain fatty acid uptake, including CD36, FABP family members, and SLC27A1/4. However, these genes share the same model reaction and therefore cannot be distinguished as independent functional dependencies from the present analysis. Moreover, the elevated uptake signal was confined to one Gleason 3+3 position (P1.2), indicating a localized model prediction rather than a consistent metabolic feature of the malignant regions. Removal of exogenous lipids reduced predicted biomass by 8–17%, supporting a model-level contribution of extracellular lipid availability under the specified nutrient conditions. Nevertheless, this result does not establish transporter-specific essentiality or cancer-selective sensitivity. The analysis therefore identifies fatty acid uptake as a candidate region-specific metabolic dependency that warrants testing through transporter perturbation, lipid uptake measurements, and growth assays under controlled nutrient conditions.

### 3.5 Independent datasets highlight the limits of transcriptome-derived target prioritization

To assess the biological context of the flux-based predictions, we compared prioritized genes with independent transcriptomic and gene-essentiality datasets. These comparisons revealed limited concordance. Although the model prioritized CD36-associated fatty acid uptake, CD36 transcript abundance was lower in bulk TCGA prostate tumors than in normal tissue (log2FC −1.40), as was that of SLC7A5 (−0.99), and DepMap data provided little evidence of selective dependence on the prioritized uptake genes: only CD36 was essential in any prostate cancer line (22Rv1, Chronos −0.51). Conversely, FASN was transcriptionally elevated in tumors (log2FC +1.65, adjusted p = 3.3 × 10⁻¹¹), as were HK2 (+0.57, adjusted p = 1.9 × 10⁻⁴) and, only slightly, ACLY (+0.38), yet none of these showed increased flux in the modeled malignant positions; HMGCR and ACO2 did not differ (Figure 5a; Table 3). Genes that were essential in DepMap, such as HMGCR (mean Chronos −1.25 in prostate cancer lines), PKM, ACO2 and SCD, were equally essential in non-cancerous prostate lines (HMGCR −1.24; Figure 5b). These differences emphasize that transcript abundance, predicted reaction flux, and genetic essentiality represent distinct biological quantities and should not be interpreted interchangeably.

Importantly, bulk tumor measurements may obscure spatially localized metabolic programs, while single-gene perturbations may not capture dependencies involving redundant transport systems. Nevertheless, these explanations remain hypotheses rather than demonstrated resolutions of the observed discrepancies. The external comparisons therefore serve as an assessment of the current model’s interpretive limits and identify specific predictions requiring direct experimental testing.

### 3.6 Energy metabolism predictions expose sensitivity to model constraints

LDH carried 2.86–18.08 a.u. in every position, lactate secretion equaled LDH flux, and the only other lactate-producing reaction carried ≤ 2.3 × 10−4 a.u. (Figure 6a, b). At 95% and 100% of maximal biomass, the loopless LDH minimum was ≤ 0 in all positions, so growth does not require LDH in these models. This does not demonstrate the absence of aerobic glycolysis in tissue. In every state, normal included, 86–92% of the glucose carbon taken up left as lactate, glycolysis supplied 82–87% of ATP and ATP synthase only 1.3–5.4% (Figure 6c; Table 4). This partition reflects how multi-subunit respiratory complexes are bounded in E-Flux and by flux minimization; even with AND = arithmetic mean, the ATP-synthase share stayed at 4–5%. These numbers should not be read as evidence for or against a Warburg phenotype, which needs isotope tracing [50], [51]. Cancer positions did not show higher glucose uptake or lactate secretion than normal positions **(Table 4)**.

**Figure 6.**
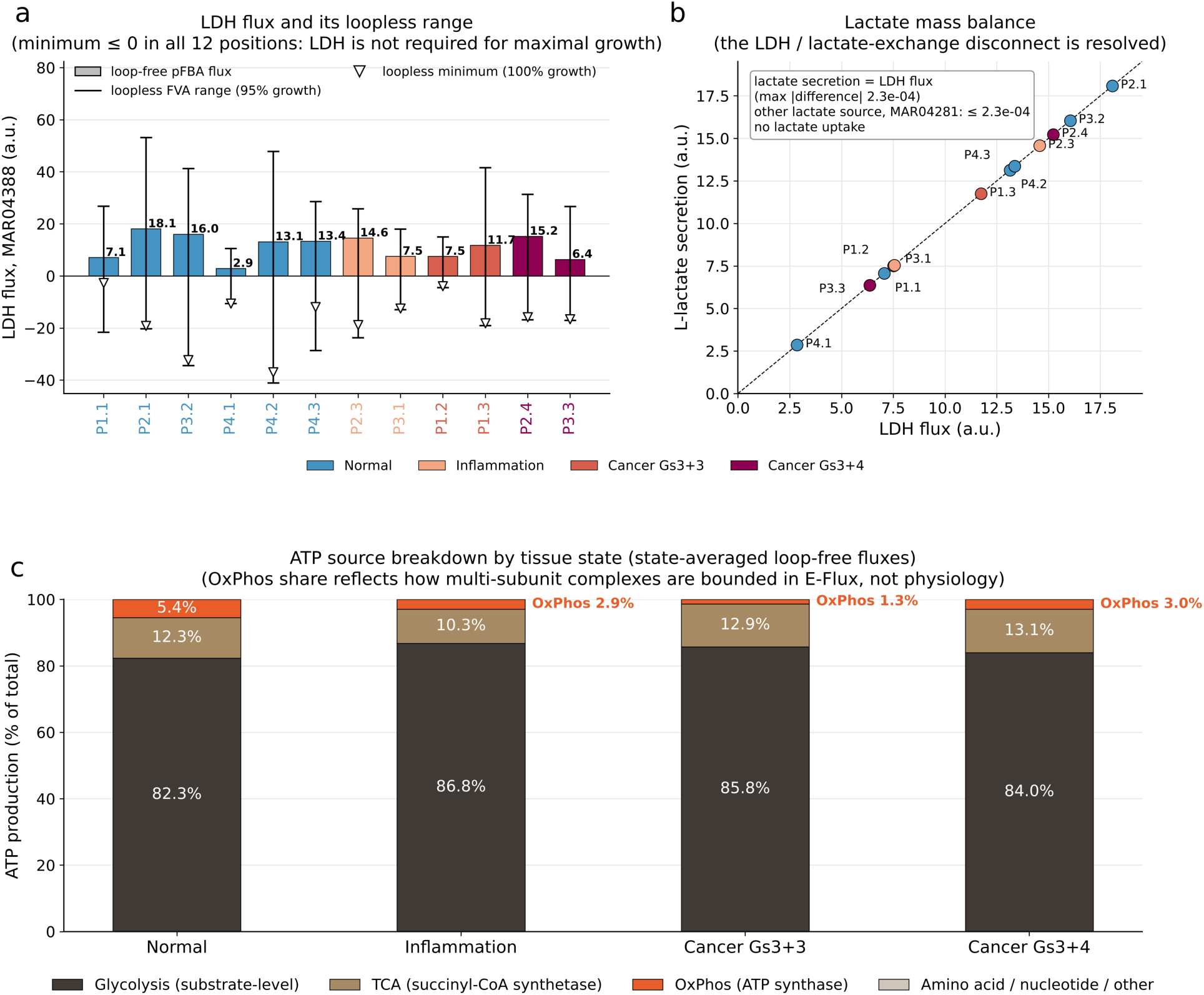
Energy metabolism in the loop-free models. **(a)** LDH flux (MAR04388) in the loop-free pFBA solution of each position (bars), its loopless FVA range at 95% of maximal biomass (whiskers) and its loopless minimum at 100% (triangles). **(b)** Lactate mass balance: lactate secretion versus LDH flux; the dashed line is identity. **(c)** ATP production by source, averaged within tissue states. The OxPhos share reflects how multi-subunit complexes are bounded in E-Flux and is not a physiological estimate.

**Table 4.**
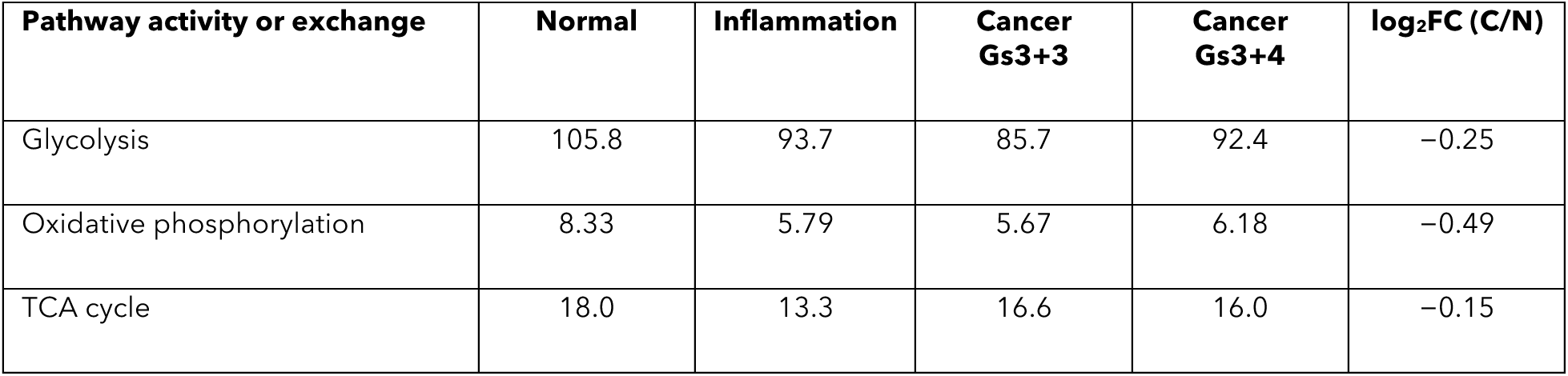
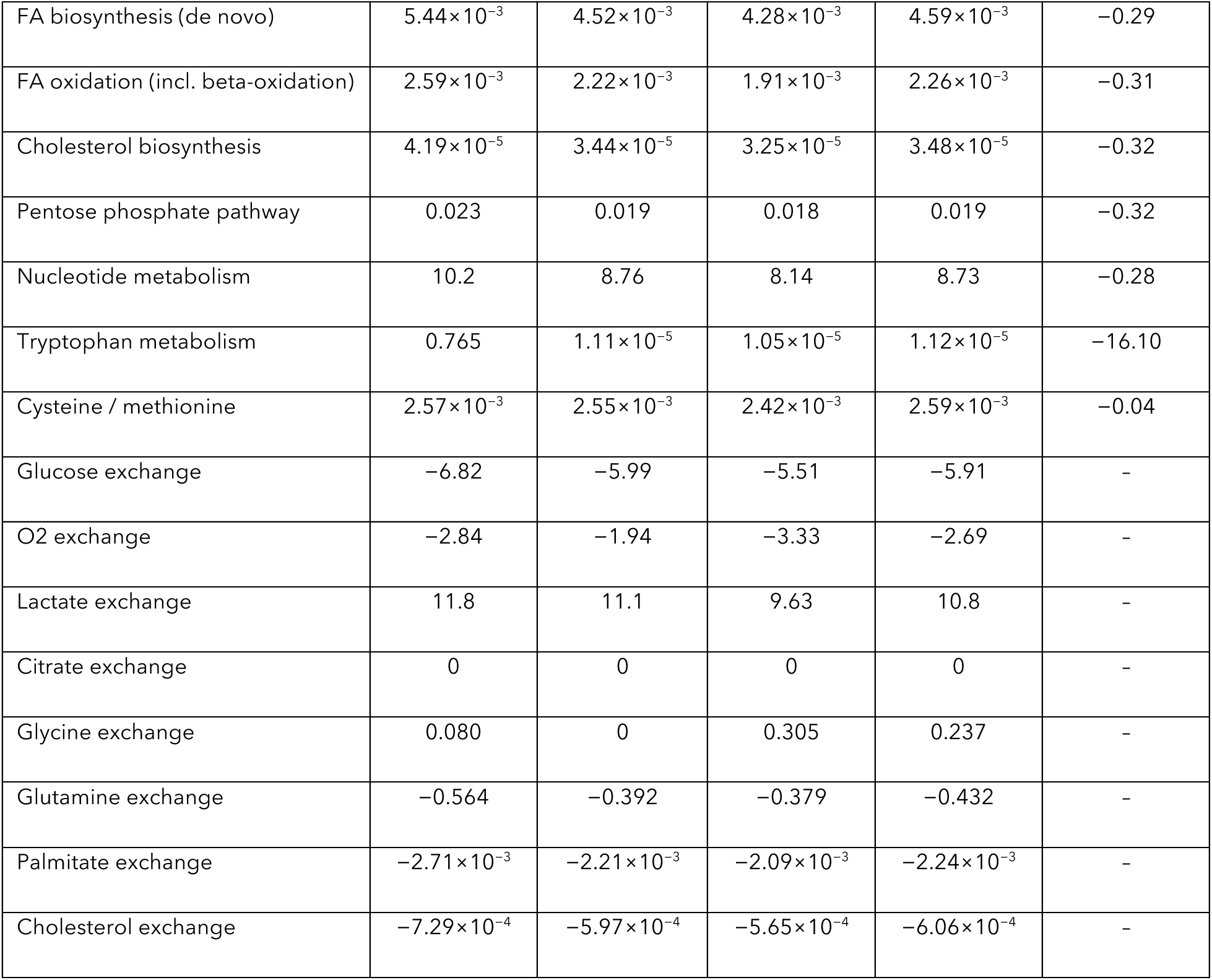
State-averaged pathway activities (Σ|v|, a.u.) and exchange fluxes (a.u.; negative = uptake, positive = secretion) of the loop-free models. log_2_FC (C/N) compares the mean of the four cancer positions with the mean of the six normal positions; tryptophan metabolism is active in three normal positions only.

| Pathway activity or exchange | Normal | Inflammation | Cancer Gs3+3 | Cancer Gs3+4 | $\log_2FC$ (C/N) |
| --- | --- | --- | --- | --- | --- |
| Glycolysis | 105.8 | 93.7 | 85.7 | 92.4 | -0.25 |
| Oxidative phosphorylation | 8.33 | 5.79 | 5.67 | 6.18 | -0.49 |
| TCA cycle | 18.0 | 13.3 | 16.6 | 16.0 | -0.15 |
| FA biosynthesis (de novo) | $5.44 \times 10^{-3}$ | $4.52 \times 10^{-3}$ | $4.28 \times 10^{-3}$ | $4.59 \times 10^{-3}$ | -0.29 |
| FA oxidation (incl. beta-oxidation) | $2.59 \times 10^{-3}$ | $2.22 \times 10^{-3}$ | $1.91 \times 10^{-3}$ | $2.26 \times 10^{-3}$ | -0.31 |
| Cholesterol biosynthesis | $4.19 \times 10^{-5}$ | $3.44 \times 10^{-5}$ | $3.25 \times 10^{-5}$ | $3.48 \times 10^{-5}$ | -0.32 |
| Pentose phosphate pathway | 0.023 | 0.019 | 0.018 | 0.019 | -0.32 |
| Nucleotide metabolism | 10.2 | 8.76 | 8.14 | 8.73 | -0.28 |
| Tryptophan metabolism | 0.765 | $1.11 \times 10^{-5}$ | $1.05 \times 10^{-5}$ | $1.12 \times 10^{-5}$ | -16.10 |
| Cysteine / methionine | $2.57 \times 10^{-3}$ | $2.55 \times 10^{-3}$ | $2.42 \times 10^{-3}$ | $2.59 \times 10^{-3}$ | -0.04 |
| Glucose exchange | -6.82 | -5.99 | -5.51 | -5.91 | - |
| O2 exchange | -2.84 | -1.94 | -3.33 | -2.69 | - |
| Lactate exchange | 11.8 | 11.1 | 9.63 | 10.8 | - |
| Citrate exchange | 0 | 0 | 0 | 0 | - |
| Glycine exchange | 0.080 | 0 | 0.305 | 0.237 | - |
| Glutamine exchange | -0.564 | -0.392 | -0.379 | -0.432 | - |
| Palmitate exchange | $-2.71 \times 10^{-3}$ | $-2.21 \times 10^{-3}$ | $-2.09 \times 10^{-3}$ | $-2.24 \times 10^{-3}$ | - |
| Cholesterol exchange | $-7.29 \times 10^{-4}$ | $-5.97 \times 10^{-4}$ | $-5.65 \times 10^{-4}$ | $-6.06 \times 10^{-4}$ | - |

These findings illustrate an important distinction between feasible metabolic flux solutions and physiologically validated metabolic states. Although the revised workflow eliminates internal cycles, the predicted partitioning of ATP production remains sensitive to expression-derived capacity constraints, nutrient availability, and the optimization objective. Consequently, the present models cannot establish the relative contributions of glycolysis and oxidative phosphorylation in prostate tissue. Experimental measurements of oxygen consumption, extracellular acidification, and isotope-resolved carbon fluxes would be necessary to evaluate these predictions quantitatively.

### 3.7 Baseline expression-constrained models do not reproduce zinc-dependent citrate regulation

ACO2 mRNA was detected in only one position (P2.1), most likely a dropout of the 2018 platform, and IDH3A was near zero (Figure 7c). SLC39A1 (ZIP1) was detected in all states, with the highest mean in Gleason 3+3 positions, opposite to the reported loss in malignant epithelium [10]; region means, however, mix cell types. In TCGA-PRAD, ACO2 was not higher in tumors (median ratio 0.88, p = 0.079; Figure 7b). In the loop-free models, citrate synthase, ACO2, IDH3 and ACLY carried no flux in any position, and Human-GEM provides no plasma-membrane route for citrate secretion (Figure 7a). Citrate synthase was optional, not blocked (loopless maximum 1.5–7.1 a.u.), so the citrate branch of the TCA cycle is not needed for growth.

**Figure 7.**
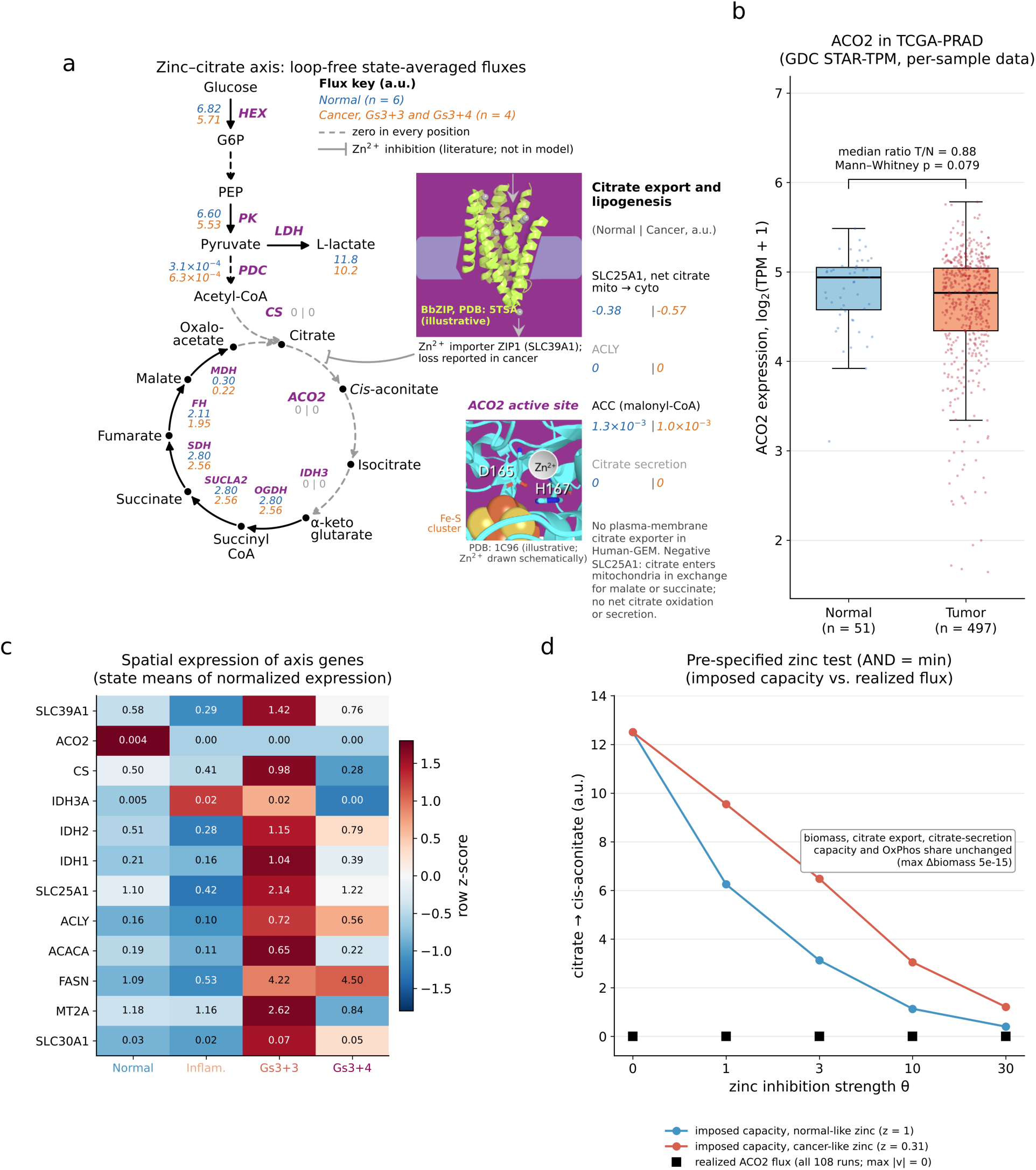
The zinc–citrate–aconitase axis under baseline assumptions. **(a)** Glycolysis and the TCA cycle with loop-free state-averaged fluxes (blue, normal, n = 6; orange, cancer Gs3+3 and Gs3+4, n = 4; a.u.). Dashed gray arrows are reactions with zero flux in every position; dashed black arrows are multi-step conversions. Cytosolic citrate reactions are listed on the right; SLC25A1 values are net citrate flux from mitochondria to cytosol, and negative values indicate citrate entry into mitochondria in exchange for malate or succinate, without net citrate oxidation or secretion. ZIP1-mediated zinc uptake and zinc inhibition of m-aconitase (ACO2) are drawn as reported in the literature [9]; this activity-level regulation is not encoded in the expression-derived bounds. The structure insets are illustrative: BbZIP from Bordetella bronchiseptica, a ZIP-family homologue of human ZIP1 (PDB 5TSA), and the active site of bovine m-aconitase in complex with citrate, showing the [4Fe-4S] cluster and catalytic residues D165 and H167 (PDB 1C96) [52]. The Zn²⁺ ion is drawn schematically and is not part of the crystal structure. **(b)** ACO2 expression in TCGA-PRAD (GDC STAR-TPM, per-sample data). **(c)** Spatial expression of axis genes (state means; color, row z-score). **(d)** Pre-specified zinc test (AND = minimum): imposed capacity of the ACO2 citrate → cis-aconitate direction (lines) and realized flux (squares; zero in all runs) as a function of inhibition strength θ.

In the pre-specified test, the imposed ACO2 capacity fell from 12.5 to 0.40 a.u. as inhibition increased, yet in all 216 simulations ACO2 and citrate synthase flux stayed at zero. Between zinc levels, biomass and citrate-secretion capacity changed by less than 10⁻¹⁰ and the ATP-synthase share by less than 10⁻⁶ (Figure 7d). Net SLC25A1 flux was unchanged except in one position under the AND = mean rule, where it shifted by up to 0.05 a.u. in both directions, consistent with alternative optima rather than a zinc effect. By the pre-specified rule, the zinc hypothesis can be neither supported nor refuted under the baseline assumptions, because the ACO2 capacity never limits flux. The baseline models therefore do not reproduce zinc-dependent citrate regulation for two identifiable reasons: the citrate branch of the TCA cycle is not required for growth, and the expression-derived ACO2 capacity is never binding.

### 3.8 Constraint-based stress testing identifies conditions permitting zinc-sensitive citrate redistribution

The absence of a zinc response under the baseline conditions prompted a post hoc analysis of the model constraints that prevent it (Supplementary Note S2; Tables S17–S20). Three constraints were identified. First, the baseline models do not require citrate oxidation. When uptake of medium carbon sources was capped in proportion to their concentrations, the models switched to oxidative metabolism: the ATP-synthase share rose from 1.1–12.3% to 22–94%, and citrate synthase became active in 5, 9 and 12 of 12 positions at the three levels of limitation (Figure 8a). Second, the network contains a route that bypasses mitochondrial aconitase. Whenever citrate was oxidized and m-aconitase was available, the parsimonious solutions routed all aconitase flux through the cytosol: citrate export by SLC25A1, ACO1, and return of isocitrate to the mitochondrion. Inhibiting m-aconitase alone therefore changed nothing.

**Figure 8.**
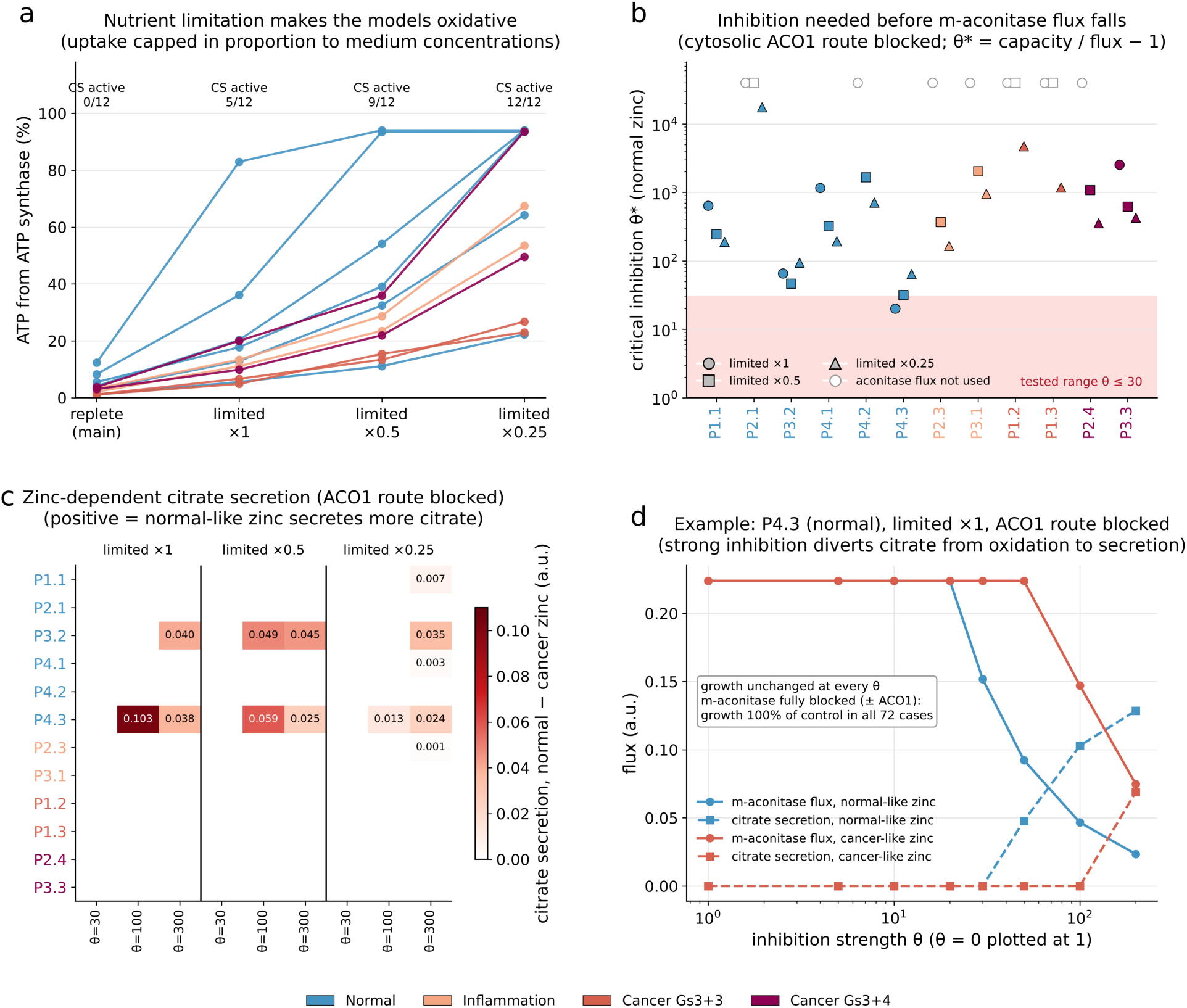
Constraint-dependent emergence of zinc-sensitive citrate fluxes in post hoc simulations. **(a)** Share of ATP from ATP synthase in the main (replete) medium and with uptake of medium carbon sources capped in proportion to their RPMI-1640 concentrations (×1, ×0.5 and ×0.25 of the unconstrained glucose uptake); numbers give the positions with active citrate synthase. **(b)** Critical inhibition strength θ* at which m-aconitase flux starts to fall, with the cytosolic ACO1 route blocked; open symbols, aconitase flux not used. The shaded band is the range tested in the pre-specified test. **(c)** Difference in citrate secretion between normal-like (z = 1) and cancer-like (z = 0.31) zinc, ACO1 route blocked, for each position, limitation level and inhibition strength. **(d)** P4.3 under limitation ×1 with the ACO1 route blocked: m-aconitase flux (solid) and citrate secretion (dashed) versus θ for normal-like and cancer-like zinc. Growth was unchanged in every simulation.

Third, with this route blocked, m-aconitase flux fell only when inhibition exceeded a critical strength θ* that ranged from 20 to 17,608 (median 398) across positions and limitation levels; it was within the originally tested range (θ ≤ 30) only in P4.3 (Figure 8b). Beyond θ*, citrate was diverted to export and secretion. Across the full grid, normal-like zinc lowered m-aconitase flux and increased citrate secretion relative to cancer-like zinc in 5 of 12 positions (P1.1, P2.3, P3.2, P4.1 and P4.3; four normal and one inflamed; Figure 8c). This direction is consistent with the mechanism proposed by Costello and Franklin [9], but it appeared only with near-complete inhibition (at least 97% in P4.3 [Figure 8d] and 99% or more elsewhere), and no position showed the opposite direction. Growth was unchanged in all 648 simulations; even complete blockade of m-aconitase, with or without ACO1, left growth at 100% of control in all 72 position–medium–route combinations, because the TCA cycle downstream of α-ketoglutarate was supplied by amino-acid catabolism. The zinc-sensitive positions were those with the lowest aconitase capacity, which was imputed from citrate synthase expression, so their ranking is not independent evidence.

The absence of a zinc response under the baseline conditions prompted a post hoc investigation of the metabolic constraints required for zinc-sensitive citrate redistribution. This analysis identified three necessary conditions within the tested model configurations: sufficient oxidative demand to activate citrate metabolism, elimination of the cytosolic aconitase bypass, and strong inhibition of mitochondrial aconitase. Under these conditions, normal-like zinc levels increased predicted citrate secretion relative to cancer-like zinc in five of twelve positions. However, the required inhibition was generally near complete, and the response did not alter predicted biomass production.

These simulations should not be interpreted as quantitative predictions of physiological zinc sensitivity. Rather, they identify network-level conditions under which the proposed zinc–aconitase mechanism becomes observable in an expression-constrained metabolic model. The results also reveal limitations of the current network representation, including alternative aconitase routes, uncertain enzyme capacities, and the absence of an explicitly represented plasma-membrane citrate secretion pathway. Direct measurements of aconitase activity, intracellular zinc, citrate secretion, and isotope-resolved TCA cycle fluxes are needed to determine whether the predicted conditions occur in prostate epithelial cells.

## 4. Discussion

The central finding of this study is that the interpretation of spatially resolved genome-scale metabolic models depends critically on the feasibility and implementation of the underlying flux calculations. Using prostate cancer spatial transcriptomics as a case study, we demonstrate that unconstrained nutrient uptake, inconsistent reaction bounds, and thermodynamically infeasible internal cycles can substantially distort predicted metabolic flux distributions. A revised workflow incorporating defined nutrient constraints, cycle certification, and loopless flux variability analysis produces mathematically admissible solutions with explicitly characterized flux uncertainty. This computational improvement does not, by itself, establish the physiological accuracy of the predicted fluxes. Instead, it provides a more defensible basis for identifying metabolic hypotheses and determining which biological measurements are necessary to evaluate them. The observed spatial differences localized fatty acid uptake signal, and conditional zinc-dependent citrate response illustrate both the potential of this approach and the limitations of inferring metabolism from transcript abundance alone.

### 4.1 Spatial and within-grade metabolic heterogeneity

With certified loop-free models, predicted metabolic similarity in this tumor tracked tissue region more closely than histological grade, and positions sharing a diagnosis could be as different as any two positions. The clearest case was the pair of Gleason 3+3 positions from the same region: one (P1.2) combined the highest AR activity with fatty-acid uptake and retinol-metabolism flux that no other position carried, whereas the other resembled the surrounding tissue. This is consistent with evidence that prostate tumors contain molecularly distinct foci not captured by Gleason grade, and that benign-appearing tissue can share alterations with nearby cancer[3], [4]. If confirmed, it would suggest that metabolic therapies, and biomarkers to select patients for them, need to account for regional variation within a tumor rather than assuming one metabolic phenotype per grade. These observations come from one patient, and regions may share technical as well as biological effects. They are hypotheses to be tested in multi-patient spatial cohorts with cell-type deconvolution.

### 4.2 Translational hypotheses: fatty-acid uptake rather than de novo synthesis

The models met lipid demand by uptake and nominated one fatty-acid uptake reaction, shared by CD36, FABPs and SLC27A1/4, and one monocarboxylate-transport reaction as cancer-elevated. FASN and ACLY carried no flux in any position, even when serum lipids were removed, HMGCR carried almost none unless serum lipids were removed, and SLC25A1 ranked only 52nd. This agrees with evidence that prostate cancer cells take up extracellular fatty acids avidly and that blocking CD36 slows tumor growth in preclinical models [15], with FABP5-dependent lipid signaling in metastasis [53], and with the preference of mammalian cells for exogenous lipids [14], [54]. It qualifies the common view of de novo lipogenesis as the main lipid vulnerability: FASN transcripts are elevated in tumors [14] and FASN inhibitors are in clinical development [55], but elevated expression does not imply that FASN flux limits growth when lipids are available. HMGCR is essential in culture but equally in non-cancerous lines, which tempers the case for statins as a cancer-selective intervention [56]. Inhibitors of SLC25A1 [57] and SLC7A5 [58] exist, but our models give them little flux-based support. Cells can also make lipogenic acetyl-CoA from glutamine by reductive carboxylation, bypassing SLC25A1 [59], [60]. In our solutions cytosolic IDH1 ran only in the oxidative direction (≤ 1.2 a.u.), but expression-constrained FBA cannot quantify this route; ¹³C-glutamine tracing is needed.

### 4.3 Zinc control of citrate fate: what the models show and what they need

The zinc–citrate–aconitase axis is a defining feature of prostate metabolism [9]. Zinc inhibits m-aconitase through metal binding [8], and the evidence for its loss in cancer comes from tissue zinc, ZIP1 protein and enzyme activities [10], [11], none of which is encoded in transcript abundance. Under the main assumptions, a pre-specified test therefore changed nothing. The post hoc analysis explains why and shows when the expected behavior appears. In the models, zinc diverted citrate from oxidation to secretion only when three conditions held together: oxidative demand (here produced by nutrient limitation), no cytosolic aconitase route around the inhibited mitochondrial enzyme, and near-complete inhibition (≥ 97–99%). In regions that met them, normal-like zinc produced the citrate-secreting phenotype and cancer-like zinc did not, the direction proposed by Costello and Franklin.

Two further results bear on the biology. First, m-aconitase was dispensable for growth in every region and condition, because the TCA cycle downstream of α-ketoglutarate was fed by amino-acid catabolism. A zinc-truncated cycle is therefore metabolically viable, in line with the secretory function of normal prostate epithelium. Second, restoring citrate oxidation was not needed for growth in cancer regions either, so the benefit of zinc loss to tumor cells is unlikely to be a simple growth requirement. These results lead to testable predictions. Zinc truncation of the cycle in prostate epithelium should coincide with limited cytosolic aconitase activity or isocitrate return, and with a high glutamine contribution to TCA intermediates downstream of α-ketoglutarate; both can be measured by enzyme assays and 13C tracing [59], [60]. The thresholds themselves depend on aconitase capacities imputed from citrate synthase expression and on the chosen nutrient limits, so they mark conditions rather than tissue values.

The same reasoning applies to energy metabolism. LDH dispensability and the glycolytic ATP partition of the main analysis reflect unlimited carbon supply; under nutrient limitation the same models become oxidative (Figure 8a), so they cannot settle whether prostate tumors are glycolytic. Likewise, AR activity varied strongly between regions but did not explain their flux differences.

### 4.4 Limitations

First, all data come from one patient; the twelve positions are not biological replicates, and regions may share section-level technical effects. Second, region-level averaging mixes epithelial, stromal and immune cells; cell-type deconvolution or spot-level modeling with higher-resolution platforms may change the results. Third, E-Flux fluxes are relative capacities in arbitrary units under an assumed biomass objective and a culture-like medium, and post-translational regulation, kinetics and metal-dependent inhibition are not represented. Fourth, 5.3% of loopless FVA bounds are conservative; conclusions that depend on them use only the existence of loop-free fluxes. Fifth, the target ranking depends on the functional form of the score and its leading signal on a single position. Sixth, the analysis of conditions for zinc control was designed after the pre-specified test and depends on aconitase capacities imputed from citrate synthase expression and on assumed nutrient limits. Seventh, no new experiments were performed; TCGA and DepMap test different properties, and isotope tracing, enzyme assays and perturbations in prostate cells or organoids remain essential.

### 4.5 Experimental validation priorities and future applications

The present study identifies several priorities for extending spatially resolved metabolic modeling beyond computational feasibility. First, independent spatial transcriptomic datasets from multiple prostate cancer patients are needed to determine whether the predicted regional differences persist across individuals and histological grades. Cell-type deconvolution or epithelial-enriched spatial profiling would help distinguish tumor-intrinsic metabolic variation from differences in tissue composition.

Second, the predicted contribution of extracellular fatty acids should be evaluated using lipid uptake assays, isotope-labeled fatty acid tracing, and targeted inhibition of candidate transport pathways. Because multiple genes map to the same uptake reaction, combinatorial perturbations may be necessary to resolve functional redundancy. These measurements would distinguish predicted reaction activity from experimentally demonstrated metabolic dependence.

Third, the zinc–citrate–aconitase axis requires measurements that cannot be inferred reliably from transcript abundance alone, including intracellular zinc concentrations, mitochondrial and cytosolic aconitase activities, and citrate production and secretion. Isotope tracing under defined nutrient conditions would help determine whether alternative aconitase routes and downstream anaplerotic pathways operate as predicted.

Finally, benchmarking model-predicted fluxes against measured exchange rates and isotope-resolved intracellular fluxes would establish whether loop-free modeling improves physiological predictive accuracy relative to conventional approaches. These experiments represent the necessary next step in translating computationally feasible spatial metabolic models into validated tools for metabolic intervention.

## 5. Conclusions

We demonstrate that model implementation choices and thermodynamically infeasible cycles can substantially alter spatially resolved genome-scale metabolic flux predictions and downstream target prioritization. Using twelve annotated regions from a single prostate tumor, we established a workflow combining expression-constrained Human-GEM models, defined nutrient conditions, explicit internal-cycle certification, and loopless flux variability analysis. All flux solutions obtained with the revised workflow were certified free of internal cycles, with 94.7% of 17,592 flux variability bounds determined exactly.

The resulting models generated exploratory hypotheses concerning spatial metabolic heterogeneity, localized fatty acid uptake, and the metabolic conditions required for zinc-sensitive citrate redistribution. These findings remain model-dependent and require independent biological validation. Importantly, the study demonstrates that eliminating infeasible flux cycles is a necessary quality-control step but is not sufficient to establish physiological accuracy.

The broader contribution is a reproducible framework for distinguishing computationally admissible flux predictions from biologically validated metabolic mechanisms. By identifying both model artifacts and specific experimental validation requirements, this approach provides a foundation for more reliable integration of spatial transcriptomics with genome-scale metabolic modeling.

## Associated Content

### Supporting Information

The Supporting information is compiled and available free of charge at the link to be added later.

## Code availability

All code used in this study, including the E-Flux FBA implementation on Human-GEM, pathway flux aggregation, permutation testing, drug target composite scoring, AR activity scoring, FVA analysis, and external validation pipelines against TCGA-PRAD and DepMap 24Q4, is publicly available at [GitHub repository URL to be added upon acceptance]. The repository includes the complete annotated Jupyter notebook (Zargar2026_spatial_prostate_GEM_loopless.ipynb), all supporting Python scripts, and processed output CSV files for all twelve spatial positions. Human-GEM is publicly available at https://github.com/SysBioChalmers/Human-GEM. The Berglund *et al*. (2018) spatial transcriptomics data used as input are available at https://db.cngb.org/stomics/datasets/STDS0000053/data. TCGA-PRAD expression data are accessible via the GDC API at https://portal.gdc.cancer.gov/, and DepMap 24Q4 CRISPR gene effect scores are available at https://depmap.org/portal/. All analyses were performed in Python 3.11 using COBRApy v0.29.1, SciPy v1.11[61], and DESeq2 v1.42[62], with package versions and environment specifications documented in the repository README.

## Author Information

Corresponding Author

## Author Contributions

**Mohammad Reza Zargar**: Conceptualization, methodology, software, formal analysis, data curation, investigation, visualization, writing original draft, review and editing. **Sunayana Malla**: Methodology, validation, data curation, review and editing. **Rajib Saha**: Supervision (S.M.), resources, review and editing, funding acquisition. **Ratul Chowdhury**: Conceptualization, supervision (M.R.Z.), resources, project administration, review and editing, funding acquisition.

## ACKNOWLEDGMENT

R.C. acknowledges support through the Iowa State University Startup Grant, the Building A World of Difference Faculty Fellowship, NSF 22-599, EPSCoR RII Track-1, Award Number #2242763, and NSF FDT-BioTech, Award Number #2533961. R.S. acknowledges support from the National Institutes of Health (NIH) Maximizing Investigators’ Research Award (MIRA) (5R35GM143009). The authors thank the developers of Human-GEM, COBRApy, and the Berglund *et al*[3]. spatial transcriptomics dataset for making their resources publicly available.

## Declaration of Conflicts of Interest

The authors have no conflicts of interest.

## Notes

### Competing Interest Statement

The authors have declared no competing interest.

### Summary of Updates

Title changed to Thermodynamic Quality Control Reshapes Spatially Resolved Metabolic Flux Predictions in Prostate Cancer. The manuscript has been substantially revised and reframed. The main contribution is now methodological, and the biological findings are presented as hypotheses. Methods: The modeling workflow was revised. It now uses a defined RPMI-1640 plus serum medium in place of open exchange reactions, scales reverse bounds of reversible reactions, and applies parsimonious FBA followed by CycleFreeFlux. Every flux solution is checked with a loop certificate and is certified free of thermodynamically infeasible cycles (TICs). All TIC-participating reactions of Human-GEM were enumerated, and the list is certified complete. Loopless flux variability analysis replaces standard FVA, and each bound is labeled as exact or conservative. Nine reaction identifiers used for key reactions in the previous version were corrected. Linear programs are solved with HiGHS. Results: A new section compares the initial and revised implementations. It shows that the initial implementation produced TICs in all twelve positions and inflated internal fluxes 190 to 890 fold. All downstream results (spatial heterogeneity, androgen receptor activity, target prioritization, energy metabolism) were recomputed with the revised workflow, and their values and interpretation were updated. Target prioritization now points to a region-specific fatty acid uptake hypothesis. Comparisons with TCGA-PRAD and DepMap 24Q4 were added as independent checks. The zinc-citrate-aconitase analysis was replaced. The earlier inhibition constant derived from molecular simulation was removed, and the axis is now tested with a pre-specified design. A post hoc constraint analysis then identifies the conditions under which zinc-sensitive citrate redistribution appears in the models. Figures and tables: All figures were revised or replaced. Figure 1 (flux certification and loopless FVA) and Figure 8 (conditions for zinc-sensitive citrate fluxes) are new. The zinc pathway map is now Figure 7a. Tables were updated. Discussion: Rewritten to emphasize model limitations, the single-patient design and experimental validation priorities. Author list updated.

