## Supplementary Material for "Thermodynamic Quality Control Reshapes Spatially Resolved Metabolic Flux Predictions in Prostate Cancer"

**Contents**


- Supplementary Methods S1. Enumeration of TIC-participating reactions and completeness certificate
- Supplementary Methods S2. Loop certificate
- Supplementary Methods S3. Loopless flux variability analysis
- Supplementary Methods S4. Target scoring, sensitivity analysis and lipid-depleted medium
- Supplementary Methods S5. Model construction details and statistics
- Supplementary Note S1. Pre-specified test of the zinc–aconitase hypothesis
- Supplementary Note S2. Post hoc analysis: conditions under which zinc controls citrate fate
- Supplementary Figures S1–S2
- Supplementary Tables S1–S20 (legends here; data in Supplementary_Tables_S1-S20.xlsx)


**Supplementary Methods S1. Enumeration of TIC-participating reactions and completeness certificate**

Let S be the stoichiometric matrix of the model (12,934 reactions) and let all exchange reactions and the biomass reaction be fixed at zero, so that only internal steady states remain. Every internal reaction i was confined to a unit box: $l_i = -1$ if the reaction is reversible in Human-GEM and 0 otherwise, and $u_i = 1$ if forward flux is allowed and 0 otherwise. A reaction direction is TIC-participating if some flux vector v with $Sv = 0$ and $l \leq v \leq u$ carries non-zero flux ($|v_i| > 10^{-7}$) in that direction, because any such v is a sum of internal cycles.

To find these directions efficiently, 30 linear programs with random objective coefficients (uniform magnitudes, random signs) were solved first. Every direction with non-zero flux in any of their optima was marked TIC-participating. Each remaining direction was then tested individually by maximizing (forward) or minimizing (reverse) its flux. This yielded 2,273 TIC-participating reactions (Table S4), of which 1,673 are annotated to the Transport reactions subsystem and 1,739 have a GPR rule.

Because a direction could in principle be missed through numerical tolerance, completeness was certified separately for forward and reverse directions. For the set J of forward directions not in the list, we solved max $\Sigma_{j \in J} t_j$ subject to $Sv = 0$, $l \leq v \leq u$ (boundaries closed), $t_j \leq v_j$ and $0 \leq t_j \leq 1$. The reverse case used $t_j \leq -v_j$. An optimum of zero proves that no direction in J can carry cyclic flux. Both optima were zero (9,099 forward and 2,796 reverse directions tested), so the list is complete.

**Supplementary Methods S2. Loop certificate**

For a flux vector v, let A be the set of internal reactions with $|v_i| > 10^{-9}$, and $s_i = \text{sign}(v_i)$. The certificate is the linear program max $\Sigma_{i \in A} s_i c_i$ subject to $S_A c = 0$, with $c_i \in [0, 1]$ if $s_i > 0$ and $c_i \in [-1, 0]$ if $s_i < 0$. Any feasible c is an internal cycle that uses only the active reactions in their current directions (a sign-conformal cycle). By the loop law [1], [2], v is free of thermodynamically infeasible cycles if and only if no such non-zero c exists, that is, if and only if the optimum is zero. A positive optimum returns the cycle itself; Table S9 lists one cycle of 76 reactions found in the P1.1 solution of the initial implementation. The certificate was applied to every solution reported in the paper (Table S10) and to every flux vector backing a loopless FVA extreme.

Fluxes carried by cycles in the solutions of the initial implementation (Table S8) were quantified by applying CycleFreeFlux [3] with the biomass flux and all exchange fluxes fixed. This returns the loop-free flux vector closest to the original in the L1 norm, and the reduction in internal $\Sigma|v|$ is the flux that had been carried by cycles.

**Supplementary Methods S3. Loopless flux variability analysis**

Loopless FVA asks for the minimum and maximum flux of a reaction over all loop-free steady states that achieve at least a fraction of maximal biomass (95% here; 100% for the LDH analysis). Solving it exactly requires a mixed-integer program over the whole network [1], which was impractical for 733 reactions in 12 positions. We therefore obtained each bound by the first applicable route below and recorded which route was used (Table S5).

**(i) Exact (acyclic).** If the reaction direction is not TIC-participating (Supplementary Methods S1), the standard LP bound is the loopless bound. Any optimal flux vector can be decomposed into a loop-free part plus sign-conformal internal cycles; a direction that belongs to no cycle has the same flux in the loop-free part, which is itself feasible, so the LP optimum is attained by a loop-free vector.

**(ii) Exact (LP optimum certified loop-free).** Otherwise, the LP optimum was passed through CycleFreeFlux with biomass fixed. If the reaction kept its optimal value, the LP bound is attained by a certified loop-free vector and is exact.

**(iii) Loop-breaking.** Otherwise, CycleFreeFlux was run a second time with the target reaction also fixed; reactions that carry flux only in this second solution belong to cycles through the target. Their current directions were blocked, and the reaction was re-optimized. This was repeated (up to 15 times) until a loop-free optimum was found [3]. If it reached the LP bound, the bound is exact. Otherwise it is conservative: the reported value is attained by a certified loop-free vector, the true loopless optimum lies between this value and the LP bound, and both are reported.

**(iv) Exact (cycle-cut MILP).** For the reactions whose ranges are interpreted in the main text (LDH, ACO2, pyruvate kinase, citrate synthase, ACLY, PDH, and the lactate and citrate exchanges), remaining conservative bounds were refined by lazy cycle-cut generation. The LP is solved; if the certificate finds a cycle C in the optimum, binary indicators $y_k$ are added for $k \in C$, with $y_k = 1$ forbidding reaction k from carrying flux in its cycle direction, together with the cut $\Sigma_{k \in C}\, y_k \geq 1$. The resulting MILP is re-solved and the procedure repeated until the optimum is loop-free (time limit 120 s per bound). Cuts exclude only flux vectors that contain the identified cycle, so a converged optimum is the exact loopless optimum.

Over all positions, 13,620 bounds (77.4%) were exact by route (i), 908 (5.2%) by route (ii), 2,133 (12.1%) by route (iii) and 1 by route (iv); 930 (5.3%) were conservative. Of the 816 bounds of the 34 key reactions (Table S6), 756 (92.6%) were exact. The conservative ones concern mainly succinyl-CoA synthetase, the upper bound of pyruvate kinase, the ZIP1 transport reaction, IDH1/IDH2 and two LDH lower bounds. Linear programs were solved with HiGHS [4] (dual simplex for loop-breaking). Results of the fast LP engine were checked against a second, independently configured engine (maximum absolute difference $10^{-10}$).

**Supplementary Methods S4. Target scoring, sensitivity analysis and lipid-depleted medium**

For each reaction, Cancer_flux and Normal_flux are the mean |v| over the four cancer positions (P1.2, P1.3, P2.4, P3.3) and the six normal positions in the loop-free pFBA solutions. Fold change is $\log_2$FC = $\log_2$[(Cancer_flux + ε)/(Normal_flux + ε)] with $\varepsilon = 10^{-9}$. Because of this pseudocount, reactions with zero flux in every normal position have $\log_2$FC ≈ 24–29; Figure 4a marks them. ΔFVA is the mean loopless FVA span (maximum − minimum) in cancer positions minus that in normal positions, divided by the largest absolute ΔFVA over all reactions and clipped at zero. The score is Cancer_flux × max($\log_2$FC, 0) × (1 + $\Delta FVA_{norm}$).

Candidates were reactions with a GPR rule, Cancer_flux > $10^{-5}$ a.u. and $\log_2$FC > 0 (94 reactions). Gene scores were the maximum over the gene's candidate reactions. Genes were prioritized if their score was ≥ the 85th percentile of gene scores, their Cancer_flux ≥ the median and $\log_2$FC ≥ 0.5 (13 genes). Three alternatives were computed: (a) ΔFVA from standard rather than loopless FVA; (b) no ΔFVA term; and (c) an additive score, the sum of the percentile ranks of Cancer_flux, $\log_2$FC and loopless ΔFVA. Each was compared with the main ranking by Spearman correlation and by the overlap of the top 20 genes (Table S12c).

For the lipid-depleted scenario, uptake of palmitate, oleate, stearate, arachidonate and cholesterol was set to zero, while the essential fatty acids linoleate and linolenate were retained. All twelve models were then re-solved with the same pFBA and CycleFreeFlux procedure (Table S13).

**Supplementary Methods S5. Model construction details and statistics**

*Model.* Human-GEM was loaded from SBML (12,931 reactions). The prostate biomass reaction (Table S2) and two transport reactions it requires (isoleucine, extracellular → cytosol; tyrosine, cytosol → mitochondria) were added, and the generic biomass reactions MAR13082 and MAR12140 were blocked. The defined medium (Table S1) allows uptake through 60 exchange reactions with unbounded capacity (−1000); the remaining 1,600 exchanges are secretion-only.

*E-Flux.* Each GPR rule was parsed recursively: AND → minimum over subunits, OR → sum over isozymes [5]. Genes not detected in a position took that position's smallest positive expression value. Weights were scaled per position as $c_i = 1000 \cdot w_i/\max_k w_k$. Reversible reactions received $[-c_i, c_i]$ and irreversible reactions $[0, c_i]$; reactions without a GPR kept their Human-GEM bounds. The twelve resulting models are provided as SBML files.

*Spatial statistics.* Pathway activity is Σ|v| over the reactions of each subsystem, excluding exchange and biomass reactions. Subsystems with activity > $10^{-7}$ in at least one position were retained (65 subsystems; Table S14). Metabolic distance was 1 − Pearson r over all 12,934 fluxes. For region and state, the test statistic was the Mann–Whitney U comparing same-label with different-label pairwise distances. The null distribution was obtained by permuting labels among the twelve positions (10,000 permutations, seed 42), which preserves the dependence among pairs that share a position, and $p = (b + 1)/(m + 1)$ [6]. A pairwise Mann–Whitney p-value is reported for description only. Principal component analysis [7] used $\log_{10}(\text{activity} + 10^{-9})$, standardized per subsystem. Biomass was compared across the four states by the Kruskal–Wallis test [8].

*AR activity.* The score is the mean expression of AR, KLK3, KLK2, TMPRSS2, FKBP5, NKX3-1 and STEAP2 per position (Table S15b). Spearman correlations with the 60 pathway activities that vary across positions were tested two-sided, with Benjamini–Hochberg adjustment [9] (Table S15).

*External data.* TCGA-PRAD per-sample STAR-TPM values (497 primary tumors, 51 solid-tissue normals) were compared by two-sided Mann–Whitney tests, with Benjamini–Hochberg adjustment across the genes tested; $|\log_2 FC| \geq 0.5$ of median TPM with adjusted $p < 0.001$ was considered supportive. ZIP1 (SLC39A1) values were excluded because the cached column had been retrieved under an incorrect Ensembl identifier (ENSG00000158714, SLAMF8). DepMap 24Q4 Chronos gene effects were taken for five prostate adenocarcinoma lines and five non-cancerous prostate lines with CRISPR data, with essentiality defined as an effect $< -0.5$ (Table S16).

**Supplementary Note S1. Pre-specified test of the zinc–aconitase hypothesis**

The following design was written down before any zinc-scenario simulation was run. All results are reported (Tables S11, S11b).

**Hypothesis.** Normal prostate epithelium accumulates zinc, which competitively inhibits mitochondrial aconitase (citrate → cis-aconitate only), truncating citrate oxidation and favoring citrate accumulation and secretion. Malignant tissue has 62–75% less zinc, relieving the inhibition and increasing citrate oxidation [10–12].

**Literature inputs.** Inhibition is competitive with citrate and acts only on the citrate → cis-aconitate direction; cytosolic aconitase is not inhibited [10]. Relative zinc in cancer $z_C = 0.31$ (range 0.25–0.38); normal $z_N = 1$. No numeric $K_i$ is available, so the inhibition strength $\theta = [\text{Zn}]_{\text{normal}}/K_i$ was swept over {0, 1, 3, 10, 30}; $\theta = 0$ is the uninhibited control.

**Implementation.** (1) The capacity of the citrate → cis-aconitate direction of ACO2 (MAR04458) was multiplied by $1/(1 + \theta z)$, the competitive-inhibition factor in the low-citrate limit, i.e. the strongest possible effect. The reverse direction and ACO1 were unchanged. (2) ACO2 mRNA is detected in only one of twelve positions, a technical dropout because ACO2 is ubiquitously expressed. Baseline ACO2 capacity in each position was therefore set to that position's citrate synthase (CS, MAR04145) capacity, CS being the adjacent TCA step and detected in all positions; this applies to both ACO2 reactions (MAR04458, MAR04589). (3) Human-GEM has no route for citrate secretion, so a curated uniport citrate[c] → citrate[e] (no GPR) was added and treated as TIC-participating. (4) Main analysis with AND

= minimum; pre-specified sensitivity with AND = arithmetic mean of subunit expression. (5) Everything else as in the main pipeline (defined medium, symmetric bounds, pFBA → CycleFreeFlux → loop certificate, loopless FVA at 95%).

**Design.** Counterfactual: every position simulated under both zinc levels, isolating the zinc effect from expression differences. Biological scenario: normal and inflamed positions at $z_N$, cancer positions at $z_C$.

**Outcomes.** O1, ACO2 citrate → cis-aconitate flux (loop-free pFBA); O2, fraction of mitochondrial citrate oxidized by ACO2 versus exported by SLC25A1; O3, maximal loop-free citrate secretion at ≥ 95% of maximal growth (loopless FVA); O4, share of ATP from ATP synthase. Prediction under the hypothesis: lowering zinc increases O1 and O4 and decreases O3.

**Decision rule.** The hypothesis is "supported by the model" if the predicted direction holds in the biological scenario and in the counterfactual for $\theta \geq 1$ under at least the main AND rule; otherwise "not supported" or "indeterminate".

**Result.** In all 216 runs, O1 was zero (ACO2 and CS carried no flux), so O2 was undefined. O3 was 20.7 a.u. (AND = minimum) and 21.1 a.u. (AND = mean) and changed by $< 10^{-10}$ between zinc levels; O4 changed by $< 10^{-6}$; biomass changed by $< 10^{-10}$, and all loop certificates were zero. Net SLC25A1 flux was unchanged except in P1.1 under AND = mean, where it shifted by up to 0.05 a.u. in both directions across θ, consistent with alternative optima rather than a zinc effect. The imposed ACO2 capacity fell from 12.5 a.u. (θ = 0) to 0.40 a.u. (θ = 30, normal zinc) but never became binding. Verdict: indeterminate; the model cannot resolve the hypothesis.

**Supplementary Note S2. Post hoc analysis: conditions under which zinc controls citrate fate**

This analysis was designed after the pre-specified test (Note S1) returned no effect, to find out why and whether any condition reveals one. It does not change the verdict of Note S1. All runs are reported (Tables S17–S20); scripts are in analysis/zinc_conditions of the repository.

**Nutrient limitation.** In the main analysis, uptake of medium components is unbounded, so carbon is never scarce. Here, uptake of the medium carbon sources (glucose, the 20 RPMI-1640 amino acids and serum lipids) was capped in proportion to their RPMI-1640 concentrations (glucose 11.1 mM, glutamine 2.05 mM, other amino acids 0.02–1.15 mM; serum lipids, approximate values for 10% FBS, 0.003–0.06 mM). The scale factor was chosen so that the glucose cap equalled ×1, ×0.5 or ×0.25 of the glucose uptake in the unconstrained solution of the same position. Vitamins, ions, $O_2$ and water were left unbounded. Everything else (E-Flux bounds with AND = minimum, imputed ACO2 capacity, curated citrate exporter, pFBA → CycleFreeFlux → loop certificate) was as in Note S1.

**Cytosolic aconitase route.** In the parsimonious solutions, citrate oxidation used the cytosolic route: mitochondrial citrate exported by SLC25A1 (MAR02405), cytosolic aconitase ACO1 (MAR04454), and isocitrate returned to the mitochondrion (MAR06289, MAR06291) for IDH3. Because zinc inhibits m-aconitase but not cytosolic aconitase [10], each scenario was run with this route open (as in Human-GEM) or blocked (ACO1 set to zero).

**Grid.** For every position: three limitation levels × two route settings × $\theta \in \{0, 10, 30, 100, 300\}$ × relative zinc $z \in \{1, 0.31\}$ ($z = 1$ only at $\theta = 0$), i.e. 648 simulations (Table S18). The critical inhibition $\theta^*$ is the strength at which the imposed capacity equals the m-aconitase flux of the uninhibited solution: $\theta^* = \text{capacity}/\text{flux} - 1$ (normal zinc). Because the parsimonious flux is at least the minimal requirement, $\theta^*$ is a lower bound. Its validity was checked by fine sweeps in P4.3 and P3.2 (Table S20). Necessity of m-aconitase was tested by blocking its citrate → cis-aconitate direction completely, with ACO1 open or blocked, in the replete medium and at limitation ×0.5 and ×0.25 (72 cases; Table S19).

**Results.** (i) With limited uptake the models became oxidative: the ATP-synthase share rose from 1.1–12.3% (replete) to 22–94% (×0.25), and citrate synthase was active in 0, 5, 9 and 12 of 12 positions (replete, ×1, ×0.5, ×0.25; Table S17). (ii) With the cytosolic route open, all aconitase flux went through ACO1 (m-aconitase flux $\leq 2.5 \times 10^{-9}$ in all 26 cases with active citrate synthase), so m-aconitase inhibition had no effect at any $\theta$. (iii) With the route blocked, $\theta^*$ ranged from 20 to 17,608 (median 398; 26 finite values), and was $\leq 30$ only for P4.3 at ×1. (iv) Normal-like zinc lowered m-aconitase flux and raised citrate secretion relative to cancer-like zinc in P1.1, P2.3, P3.2, P4.1 and P4.3 (Table S18); the contrast appeared from $\theta = 30$–100 in P4.3 and P3.2 and only at $\theta = 300$ elsewhere, and the opposite direction never occurred. (v) Growth varied by less than $2 \times 10^{-11}$ (relative) across all 648 simulations, and complete blockade of m-aconitase left growth at 100% of control in all 72 cases.

**Interpretation.** In these models zinc controls citrate fate only when oxidative demand, absence of the cytosolic route and near-complete inhibition coincide. m-aconitase is never required for growth, because α-ketoglutarate and downstream TCA intermediates are supplied by amino-acid catabolism. The positions that respond first are those with the lowest imputed aconitase capacity; because that capacity was imputed from citrate synthase expression, the spatial ranking of zinc sensitivity is a consequence of the imputation and not independent evidence.

### Supplementary Figures

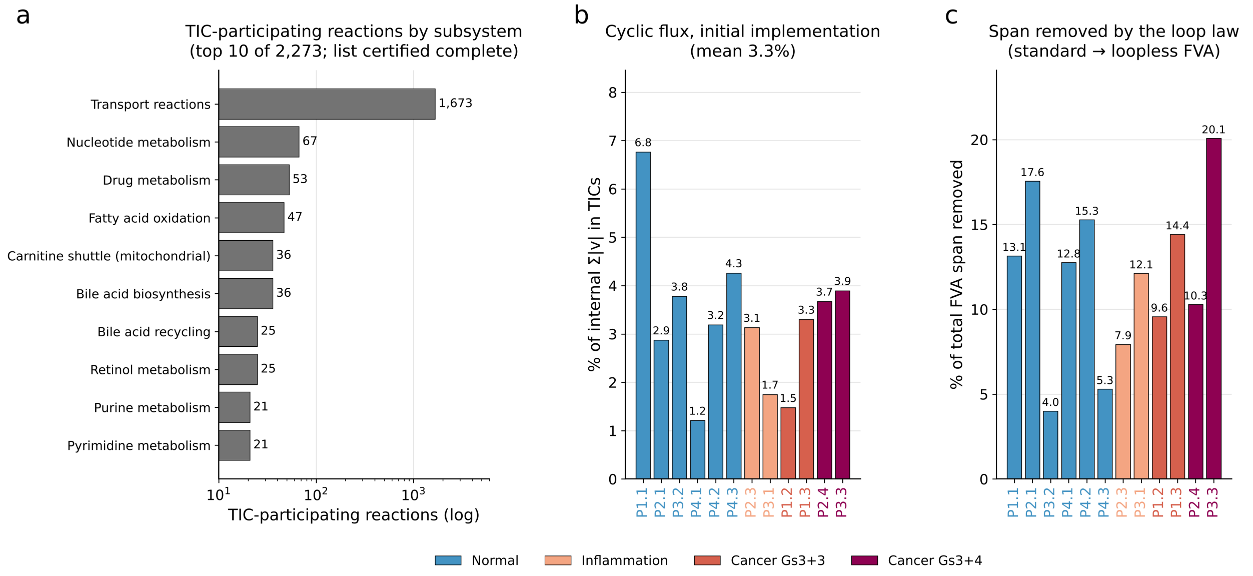


**Figure S1. Thermodynamically infeasible cycles in detail.** (a) TIC-participating reactions of Human-GEM by subsystem (top 10 of 2,273; the list is certified complete, Supplementary Methods S1). (b) Percentage of internal flux carried by TICs in the flux solutions of the initial implementation, obtained by removing cycles with biomass and exchanges fixed. (c) Percentage of the total standard FVA span removed by the loop law in each position (≥ 95% of maximal biomass; 733 reactions).

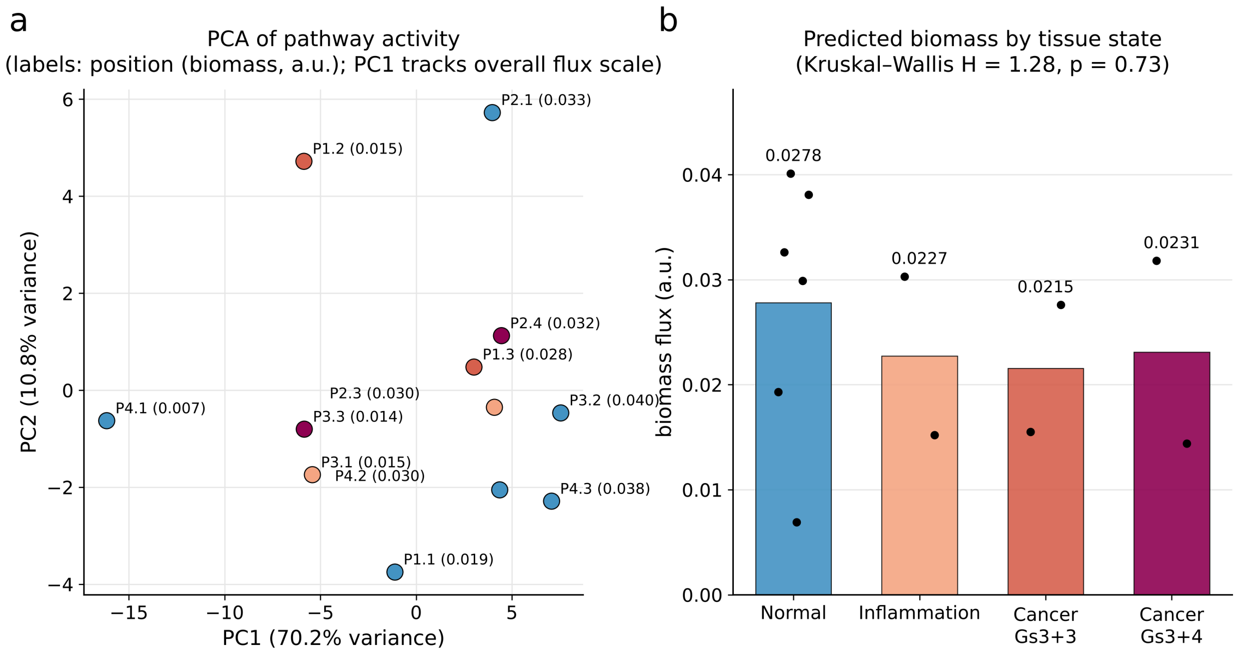


**Figure S2. Principal components and biomass.** (a) Principal component analysis of the 65 active pathway activities (log-scaled, standardized); labels give position and predicted biomass. PC1 mainly reflects overall flux scale (P4.1, lowest biomass, at one extreme) and is not interpreted as a biological axis. (b) Predicted biomass by tissue state (bars, means; points, positions); with two positions in three of the states, the test has little power.

**Supplementary Tables**

All supplementary tables are provided in the workbook Supplementary_Tables_S1-S20.xlsx, one sheet per table, with a Contents sheet. Fluxes are in arbitrary units (relative E-Flux capacities); exchange fluxes are negative for uptake and positive for secretion. Table S3 is also reproduced below because it documents the reaction-identifier corrections between the initial and revised implementations.

**Table S1.** Defined medium: the 60 uptake-permitted exchange reactions (RPMI-1640 and 10% FBS components).

**Table S2.** Composition of the prostate-cell biomass reaction.

**Table S3.** Reaction identifiers used to report key fluxes in the initial implementation that pointed to other reactions, and the correct reactions (reproduced below).

**Table S4.** The 2,273 TIC-participating reactions of Human-GEM, with the directions in which each can carry cyclic flux, subsystem, GPR status and equation.

**Table S5.** Standard and loopless FVA bounds for 733 reactions in 12 positions, with the route by which each loopless bound was obtained.

**Table S6.** Key-reaction registry: reactions used to report key enzyme and exchange fluxes, with equations.

**Table S7.** Stepwise correction of the initial setup (all solved with HiGHS): biomass, internal and exchange $\Sigma|v|$ and loop certificate per position and step.

**Table S8.** Audit of the flux solutions of the initial implementation: loop certificate, internal $\Sigma|v|$ before and after cycle removal and share of flux carried by TICs.

**Table S9.** One certified cycle (76 reactions) in the P1.1 solution of the initial implementation.

**Table S10.** Flux-solution log of the revised implementation: biomass, $\Sigma|v|$ and loop certificate after FBA, pFBA and CycleFreeFlux.

**Table S11.** Pre-specified zinc test: all 216 runs (S11) and summary by $\theta$ and zinc level (S11b).

**Table S12.** Target scoring: gene-level scores and variants (S12), reaction-level inputs for the 94 candidate reactions (S12b) and agreement of alternative formulations (S12c).

**Table S13.** State-averaged fluxes in the main and lipid-depleted media.

**Table S14.** Pathway activity (Σ|v| per Human-GEM subsystem) per position.

**Table S15.** Spearman correlations of AR activity with the 60 pathway activities (S15), and AR readout-gene expression and scores per position (S15b).

**Table S16.** TCGA-PRAD differential expression and DepMap 24Q4 essentiality for prioritized targets and reference genes.

**Table S17.** Post hoc analysis (Note S2): replete and nutrient-limited regimes with the cytosolic aconitase route open or blocked, at θ = 0 and θ = 30 for both zinc levels, for all 12 positions.

**Table S18.** Post hoc analysis: full grid of nutrient limitation × aconitase route × θ × zinc level (648 simulations).

**Table S19.** Post hoc analysis: growth with the citrate → cis-aconitate direction of m-aconitase fully blocked, with ACO1 open or blocked (72 cases).

**Table S20.** Post hoc analysis: fine sweep of θ in P4.3 and P3.2 (limitation ×1) for both zinc levels and route settings.

Table S3. Reaction-identifier corrections. Each row gives the label used in the initial implementation, the identifier that was used, what that identifier actually is in Human-GEM, and the correct reaction used in this study.

| **Label in the initial implementation** | **ID used** | **What that ID is in Human-GEM** | **Correct reaction** |
|---|---|---|---|
| SLC25A1 citrate export | MAR03964 | ATP phosphohydrolase \| ATP + H2O --> ADP + H+ + Pi | SLC25A1 citrate carrier (11 reactions, summed) |
| ACLY | MAR04934 | \| acetate + Na+ --> acetate + Na+ | MAR04149 |
| LDH | MAR04358 | ATP:pyruvate 2-O-phosphotransferase \| ADP + H+ + PEP --> ATP + pyruvate | MAR04388 |
| Citrate synthase | MAR04373 | D-glyceraldehyde-3-phosphate:NAD+ oxidoreductase (phosphorylating) \| 1,3-bisphospho-D-glycerate + H+ + NADH <=> D-glyceraldehyde 3-phosphat | MAR04145 |
| Hexokinase | MAR04319 | ATP:D-fructose 6-phosphotransferase \| ATP + fructose --> ADP + fructose-6-phosphate + H+ | MAR04394 |
| ATP synthase | MAR06921 | NADH:ubiquinone oxidoreductase \| 5.0 H+ + NADH + ubiquinone --> 4.0 H+ + NAD+ + ubiquinol | MAR06916 |
| PGK | MAR04301 | UTP:D-fructose-6-phosphate 1-phosphotransferase \| fructose-6- | MAR04368 |

<table>
<tr><td></td><td></td><td>phosphate + UTP --> fructose-1,6-bisphosphate + H+ + UDP</td><td></td></tr>
<tr><td>Pyruvate kinase</td><td>MAR04348</td><td>L-threonine ammonia-lyase (2-oxobutanoate-forming) | threonine --> 2-oxobutyrate + H+ + NH3</td><td>MAR04358</td></tr>
<tr><td>Succinyl-CoA synthetase</td><td>MAR04384</td><td>Exchange of ceramide pool | ceramide pool --></td><td>MAR04152</td></tr>
</table>
